## Supplementary material for "Isometric Spiracular Scaling in Scarab Beetles: Implications for Diffusive and Advective Oxygen Transport": Newly accepted relevant paper

**Subject:** Integrative and Comparative Biology - Decision on Manuscript ID ICB-2022-0023.R1
**Date:** Friday, June 17, 2022 at 3:17:58 PM Mountain Standard Time
**From:** Integrative and Comparative Biology
**To:** Jon Harrison
**CC:**

17-Jun-2022

Dear Prof. Harrison,

Congratulations! On the advice of the referees, your manuscript, entitled "Size-dependent scaling of stingless bee flight metabolism reveals an energetic benefit to small body size" has been accepted for publication in its current form to Integrative and Comparative Biology..

We thank you for your efforts, and we look forward to seeing your work in print. Soon, you'll receive information about page proofs. Please recognize that during that process, additional minor issues might arise and need to be addressed. We appreciate your quick attention to those.

OPTIONAL OPEN ACCESS - Integrative and Comparative Biology authors can opt, at an additional charge, to make their paper freely available online to all, including non-subscribers, immediately upon publication, as part of the Oxford Open initiative. In order to publish your manuscript, Oxford University Press requires that you complete a license agreement online. A link to the online licensing system, and instructions on how to select and complete a license, will be provided to you by the Production Editor at Oxford University Press shortly after your article is accepted. Under Step One of the form, indicate whether you wish to be published under an Open Access model and agree to pay the relevant additional charge. Applicable Open Access charges can be found at

http://www.oxfordjournals.org/icb/for_authors/index.html.

If you do not select the Open Access option, your paper will be published in the journal with standard subscription-based access, online and in hard copy, and you will not be charged. Authors who choose the Open Access publication option are also required to complete an Open Access charge form online at http://www.oxfordjournals.org/oxfordopen/forms.

Thank you for your contribution. On behalf of the Editors of Integrative and Comparative Biology, we look forward to your continued contributions to the journal.

Sincerely,
Dr. Ajna Rivera
Associate Editor, Integrative and Comparative Biology

**Title:** Size-dependent scaling of stingless bee flight metabolism reveals an energetic benefit to small body size

**Authors:** Meghan E. Duell[1], C. Jaco Klok[2], David W. Roubik[3], Jon F. Harrison[2]

**Other Affiliations:** [2]School of Life Sciences, Arizona State University, Tempe, AZ, USA; [3]Smithsonian Tropical Research Institute, Panama City, Republic of Panama.

**Synopsis:** Understanding the effect of body size on flight costs is critical for development of models of aerodynamics and animal energetics. Prior scaling studies that have shown that flight costs scale hypometrically have focused primarily on larger (> 100 mg) insects and birds, but most flying species are smaller. We studied the flight physiology of thirteen stingless bee species over a large range of body sizes (1-115 mg). Metabolic rate during hovering scaled hypermetrically (scaling slope = 2.11). Larger bees had warm thoraxes while small bees were nearly ecothermic; however, even controlling for body temperature variation, flight metabolic rate scaled hypermetrically across this clade. Despite having a lower mass-specific metabolic rate during flight, smaller bees could carry the same proportional load. Wingbeat frequency did not vary with body size, in contrast to most studies that find wingbeat frequency increases as body size decreases. Smaller stingless bees have greater relative wing surface area which may help them reduce the energy requirements needed to fly. Further, we hypothesize that the relatively larger heads of smaller species may change their body pitch in flight. Synthesizing across all flying insects, we demonstrate that the scaling of flight metabolic rate changes from hypermetric to hypometric at approximately 58 mg body mass with hypermetic scaling below (slope=1.2) and hypometric scaling (slope=0.67) above 58 mg in body mass. The reduced cost of flight likely provides selective advantages for the evolution of small body size in insects. The biphasic scaling of flight metabolic rates and wingbeat frequencies in insects supports the hypothesis that the scaling of metabolic rate is closely related to the power requirements of locomotion and cycle frequencies.

### Introduction

Understanding how body size affects animal function is one of the central themes of biology; such scaling studies have provided key syntheses of organismal function and macroecology (Sibley et al., 2012). Flight is a key trait for the evolutionary success of insects,birds, and bats being integral to resource collection (pollination), migration and defense. The scaling of flight metabolic rate with mass in insects remains a controversial issue. Studies of hovering moths and bees ranging in mass from 100-1100 mg have shown that flight metabolic rates scale hypometrically (slope less than one, indicating lower energy use per gram body mass in larger animals compared with smaller animals) with slopes of log metabolic rate on log mass of 0.63-0.77, with wing beat frequencies consistently shown to decline in larger insects (Bartholomew and Casey, 1978; Darveau et al., 2005; Suarez, 2000). In contrast, a meta-analysis by Niven and Scharlemann (2005) suggested that across all insects, flight metabolic rate scales hypermetrically (slope greater than one, indicating greater energy use per gram body mass in larger animals compared with smaller animals) with mass$^{1.1}$, and that this was due to insects below 10 mg in mass having distinctly lower flight metabolic rates than insects above this size (Casey et al., 1985). However, these authors noted that their conclusions were hampered by a dearth of studies of insect flight in the size classes across which flight costs seem to change dramatically (10-100 mg).

We aimed to determine how and why the scaling of flight metabolic rate changes with body mass among insects using stingless bees as a subset of small flying insects. The mechanisms causing hypometric scaling of metabolic rate are controversial. Supply constraint hypotheses, such as the idea that limitations on gas or nutrient transport occur due to decreasing surface-to-volume ratios or the fractal nature of delivery systems imply that consistent hypometric scaling should be observed (Cardoso et al., 2006; Nisbet et al., 2000; West et al., 2000). Conversely, the idea that the energetic costs of locomotion drive metabolic scaling suggest that a primary driver of hypometric scaling of metabolic rate in animals is the decline in cycle frequency observed in larger runners, swimmers and fliers (Biewener and Patek, 2018). Declining cycle frequency lowers cost because the rate of myosin ATPase activity increases with muscle contractile speed and contractile frequency, and likely elastic energy storage also decreases (Biewener and Patek, 2018). In most flying insects that have been examined, wing-beat frequency declines in larger species, as found in birds and bats (Casey, 1989; Darveau et al., 2005; Harrison and Roberts, 1998). Whether these trends in wingbeat frequency occur across smaller insect fliers is unknown.

We measured flight metabolic rates, wingbeat frequency, voluntary load carriage, and wing and body segment sizes in 13 species of stingless bees between 1.5-115 mg. We used stingless bees (Meliponini) for these measurements because they have a large range of body size among species, approximately 1-150 mg (Fig 1, 2A) with a well-defined molecular phylogeny

(Felsenstein, 1985; Harvey and Pagel, 1991; Ramírez et al., 2010; Rasmussen and Cameron, 2010). Some lineages, especially the genus *Melipona*, have large species ranging in body mass up to 150 mg (Camargo, 2013). Miniaturization has evolved multiple times among 33 genera (Camargo, 2013; Michener, 2001) and it is thought that ancestral Meliponines were moderately sized, perhaps 50 mg (Camargo, 2013; Camargo and Pedro, 2013; Ramirez et al., 2010; Rasmussen and Cameron, 2010). The smallest species we used, *Trigonisca buoyssoni,* was 1.5mg in size while the largest, *Melipona triplaridis*, was 115 mg ± 5 mg. Additionally, most stingless bees fill a similar ecological niche regardless of body size; all are social, living in colonies with task differentiation, and most forage on flowers resulting in a pollen and nectar diet. This diversity in the evolution of body size across the phylogeny of stingless bees and similar ecologies makes them ideal for comparative work.

### Materials & Methods

***Study sites and stingless bee collection:*** Stingless bee foragers from 11 species (*Melipona triplaridis*, *Melipona panamica*, *Scaptotrigona panamensis*, *Scaptotrigona luteipinnis*, *Trigona fulviventris*, *Trigona muzoensis*, *Tetragonisca angustula*, *Frieseomelitta nigra*, *Lestrimelitta danuncia*, *Plebeia franki*, *Plebeia frontalis*) were captured returning to nests at several locations in the Republic of Panamá. *Scaptotrigona lutipinnis*, *T. angustula*, *F. nigra*, and *T. fulviventris* were captured in Gamboa, Panamá, while *T. muzoensis*, and *P. frontalis* were collected on Barro Colorado Island. *Melipona triplaridis*, *S. panamensis*, and *L. danuncia* were collected from the property of David Roubik in Curundu, Panamá and *P. franki* and *M. panamica* were captured at the Santa Rita Arriba property of David Roubik. In each case, foragers were identified and captured as they returned to the nest from a single colony of each species. Another two species

(13 total) including *Trigonisca atomaria* and *Trigonisca buoyssoni* were collected while foraging at flowers using the canopy crane at Parque Naturál Metropolitano, Panama City, Panamá, and at Santa Rita Arriba while foraging honey water. Individuals were placed in 50 mL tubes with sugar water on cotton for food if they could not be measured within one hour of capture due to transportation time constraints. All bees were brought back to the Smithsonian Tropical Research Institute (STRI) lab in Gamboa, Panamá for measurements. We verified the identities of all species using the STRI insect collection.

***Respirometry and wingbeat frequency analysis:*** All bees were weighed on a Mettler-Toledo microbalance and placed in Fluon coated glass flight chambers for flow-through respirometry. Ambient air was pushed through silica and soda lime scrubber columns by an aquarium pump, and flow rate through the respirometry chamber was adjusted using a Sable Systems FlowBar 8 mass-flow controller (resolution ±0.1 ml/min below 100 ml $min^{-1}$; resolution 1 ml $min^{-1}$ above 100 ml $min^{-1}$ flow rate). Excurrent $CO_2$ was measured using a LiCor 6252 plumbed in the differential mode (the reference cell measured the air flowing into the chamber and the sample cell measured air flowing out of the chamber; resolution was approximately 0.2 ppm with hardware and software time-averaging of 1 sec). The system was calibrated and spanned using a $CO_2$ tank containing 1221 ppm $CO_2$ (as measured by by J. Shik with a LiCor 7000 calibrated against a certified span gas), with the zero-span reset each time the flow rate was changed and zeroed before and after each bee was measured. We used four different cylindrical glass flight chambers with volumes of 15 mL, 70 mL, 150 mL, and 550 mL; chamber sizes were adjusted to the size of the bee. We chose the smallest chamber that a species would fly consistently in. Flow rates were adjusted to chamber size so that the 95% washout time for that chamber was approximately one min or less; flow rates ranged from 150 mL min$^{-1}$ for the smallest chambers to 1000 mL min$^{-1}$ for the largest chamber used. $CO_2$ levels during flight ranged from 6 - 175 ppm, with a minimal signal-to-noise ratio of 9.5. The analog outputs of the $CO_2$ analyzer and mass flow-controller were digitized and recorded with a Sable Systems UI-2 and a computer using Expedata Pro 1.7.2 (digitization resolution was 0.5 ppm for the $CO_2$ analyzer, accounting for baseline noise, and 0.1 mL min$^{-1}$ for the mass flow-controller).

We used several methods to maintain good flight behavior including agitation of the chamber and shining a bright light above the chamber while its surroundings were kept dark. We only accepted data from bees that exhibited at least 30 sec of consistent flight behavior that was accompanied by a relatively high and consistent $CO_2$ reading measured after the time required for washout of any atmospheric $CO_2$ that may have entered the chamber when the bee was placed inside (average flight duration measured was 43 sec). After measuring $CO_2$ emission during flight, the air pump was turned off and we inserted a Sony ECM-PC60 mini electric condenser microphone to record wingbeat frequency for each bee. This was recorded and analyzed using Raven Lite 1.0 software. A subsample of 4-5 bees/species were then stimulated to fly in the same chambers and filmed with a MotionPro X high-speed video camera at 1000 fps to verify wingbeat frequency data acquired with the microphone. We used the average wingbeat frequency from three measures per individual for analysis.

***Body temperature in flight:*** After measurements of wing beat frequency, we removed bees from the chamber and placed them in a plastic gallon Ziploc bag; they continued flying within the bag until measurement of body temperature was accomplished. We used a ‘grab-and-stab’ measurement technique (Roberts and Harrison, 1999) with a Physitemp MT-29/1 hypodermic needle microprobe (29 ga, 0.025sec time constant) and a Physitemp BAT-10 thermocouple meter. To minimize thermal transfer from human to bee, we wore insulated gloves to hold the temperature microprobe, and restrained bees by pulling the plastic bag tight about them, on top of a Styrofoam board. We measured air, abdomen, and head temperatures in random order for every bee after first measuring thorax temperature; thorax temperatures were taken within 1 s of restraint and all temperatures were measured within an additional 2 s which minimized any heat transfer from the process. Later, an additional ten bees per species were killed by freezing for one hour and we measured their body segment temperatures after keeping them at room temperature, or 30 °C for 15 min (the length of time needed for body temperatures to stabilize). We used these measurements to verify that there was no heat transfer from handling or the thermocouple to bees because dead bees had body segment temperatures within 0.4 °C of the measured air temperature.

We also flew 30 *S. luteipennis* (a medium sized species) individuals at temperatures between 25 - 40 °C to establish a $Q_{10}$, which was 0.96 on average. Each bee was placed in a Fluon-lined respirometry chamber (as outlined above) that was located inside chamber with foam insulation and plastic sheeting in front to access the flight chamber. We used an ITC-306T Inkbird Temperature Controller (Inkbird, Shenzhen, China) probe, and connected the controller to a heater and fan that kept the temperature constant within the chamber (± 0.5 °C) while the bee was flying. A micro thermocouple probe was also placed inside the respirometry chamber to record temperature during flight. We flew each bee at a single air temperature and made a thermal performance curve (Fig. 2C) with data from all bees and then calculated $Q_{10}$.

***Wing morphology and load carriage:*** We removed the wings for 10 individuals/ species by removing and flattening them onto white cardstock paper with transparent tape. A digital image of each wing was taken with a 1 mm grid for calibrating measurements. All measurements were performed in ImageJ. To determine load carriage (the maximum amount of liquid that could be carried), we starved 10 bees per species for 2 hours, weighed them, then fed them 50% sucrose solution to satiation, weighed them, and encouraged them to fly as described above. If they would not fly, we continued coaxing them until they would and immediately reweighed at that time to reflect the most accurate load they could lift off and fly with.

***Phylogenetic and Statistical Analysis:*** All data for stingless bees are represented as species means ± SE (standard error) of individual measurements. The effect of body mass was tested using least squares linear regression performed on log-transformed data to obtain the metabolic rate equation $aM^b$, where a=y-intercept, M=body mass, and b= allometric scaling coefficient (Darveau et al., 2005).

We converted metabolic rates (ml $g^{-1}$ $h^{-1}$) to watts and applied corrections for various $Q_{10}$ values ($Q_{10}$ = 1 as measured in *S. luteipennis* (Fig. 2 C) and $Q_{10}$ = 2 which is more typical of hymenopterans) to compare with flight metabolic rate data from the literature (Fig 1D). We assumed RQ =1 based on available data for hymenopterans (Suarez et al., 2005) and because bees were fed solely on a diet of sucrose water while in captivity. Further analyses of wingbeat frequency, wing area, wing loading, and flight body temperature were performed using phylogenetic generalized least squares regressions (PGLS) in R on log-transformed data. A comprehensive maximum likelihood tree based on Rasmussen & Cameron (2010) and Ramirez et al. (2010) were adapted for this study by pruning unnecessary species and adding species that did not appear on the published phylogenies. Branch lengths for all tip species were then set equal to one (Garamszegi, 2014; Ramirez et al., 2010; Rasmussen and Cameron, 2010, 2007). PGLS was performed for all analyses using all statistically possible tree topologies and results were obtained using the topology with the highest likelihood (Garamszegi, 2014).

We compared the known metabolic rates of 117 flying insects by compiling literature values (Table S1). Data points were eliminated from this dataset if they did not use modern methods (i.e. flow-through or stop flow respirometry) for determining flight metabolic rates or were measured in non-standard conditions (i.e. fluctuating temperatures or humidity, dietary manipulations, carried loads, etc.). We generated all possible models for flight metabolic rate scaling, including breakpoint models of log body mass vs. log metabolic, in R using the breakpoints and lm.br packages (Priyadarshana and Sofronov, 2016). The model was unconstrained to allow discontinuous slopes on either side of breakpoints and bootstrap restart sampling between 20-60 mg body mass. This generated multiple possible piece-wise regressions that differed in slope and breakpoint. We chose the regression with the lowest error represented as MSE. We compared this piece-wise regression to the standard models with a continuous slope across body size of 0.75 and 1.0 using AIC comparisons from the breakpoints package in R.

### Results & Discussion

Flight metabolic rates scaled hypermetrically across stingless bee species, with a scaling exponent of 2.25 (Fig. 2B, Adj. $R^2$=0.66, P<0.001, $\lambda$=0.00, $P_{(0)}$ = 0.0175). This slope was not significantly affected by corrections using phylogenetically generalized least squares analysis (PGLS, Table 1). The 95% confidence limits for this slope did not include isometry (slope = 1) or the hypometric exponents found for Euglossine bees or moths (Bartholomew and Casey, 1978, 1977; Dillon and Dudley, 2004). The different scaling patterns in Euglossines and stingless bees are not due to differences in absolute cost, since at body masses at which both taxa have been measured (circa 100 mg) flight metabolic rates are very similar (Fig 2B).

Flight muscles of insects generate substantial heat, and most insects larger than 50 mg flew at thorax temperatures 5-20 °C above air temperature, while insects with body masses lower than 50 mg have high cooling rates and usually have body temperatures close to air temperature (Heinrich, 1993; May, 1981; Unwin and Corbet, 1984). Body temperatures of stingless bees demonstrated the size-related pattern expected from studies of other insects (Fig. 2C, D). Stingless bees heavier than 70 mg (*M. panamica* and *M triplaridis*) had substantially elevated body temperatures, more than 10°C above air temperature, as previously shown (Roubik and Peralta, 1983). In these two large species, thorax temperatures were the highest, as predicted by heat production in the flight muscles, and the abdomen was the coolest region (Fig. 2E). In contrast, in stingless bee species less than 20 mg, head, thorax, and abdomen temperatures were mostly uniform and only about 1-3°C above air temperature (Fig. 2D).

Might the thermal differences between large and small bees drive the hypermetric scaling of flight metabolic rates in stingless bees, with the warmer temperatures of larger bees allowing higher metabolic rates? The metabolic rates of large flying insects can increase with thorax temperature (Harrison et al., 2001; Heinrich, 1971), decrease with thorax temperature (Byrne et al., 1988; Harrison et al., 1996; Roberts and Harrison, 1999) or be independent of thorax temperature (Bartholomew and Casey, 1977; Casey, 1989, 1981; Heinrich and Casey, 1973; May, 1995), depending on the species and range of thorax temperatures examined. We found a $Q_{10}$ of 0.96 in one species of stingless bee (*Scaptotrigona luttipennis*) with no significant trend in flight metabolism over a 20° C range of thorax temperatures (Fig. 2D). If this low thermal sensitivity of flight applies to all species, then variation in body temperature will not explain variation in flight metabolism. However, for the small insects that have been measured, flight performance increases strongly over cool to moderate ranges of thoracic and air temperature, with $Q_{10}$ values for wing beat frequency, flight speed, force production, power output and metabolic rates of 1.2 – 2 (Heinrich, 1975; Henry and Harrison, 2014; Lehmann, 1999; Yurkiewicz and Smyth, 1966). To test the possibility that the lack of hypometric scaling of stingless bees was caused by thermal variation across species, we fit the thorax temperature data with a third-order polynomial (Fig. 2C), and then used these data to apply $Q_{10}$ corrections to the flight metabolic data (Fig. 2B). Using a $Q_{10}$ correction of 2, one of the highest $Q_{10}$ values found for flying insects, the scaling slope was still significantly hypermetric with a slope of 2.11 (Fig. 2 B, Table 1). Thus, the lack of hypometric scaling in flying stingless bees cannot be explained by thermal variation among these species.

The differential scaling of flight metabolic rates in stingless bees and Euglossine bees was associated with differential scaling of their wing morphology. Larger stingless bees had relatively smaller wings, as the slope of total wing area scaled with body mass with a scaling exponent of 0.56 (Fig. 2F, Table 1), significantly less than the isometric prediction of 0.67, and contrasting with the pattern for Euglossine bees, in which larger bees have relatively larger wings than predicted by isometry (Casey et al., 1985). The relatively smaller wing area in larger bees arose mostly from these wings being relatively narrower (Fig. 3, Table 1), as wing lengths scaled isometrically (Fig. 3). One possibility is that the relatively larger wings in smaller bees could create more lift per stroke, potentially reducing energetic cost and contributing to the lower flight cost per gram observed in smaller bees.

In contrast to the scaling of wing area, the masses of body segments scaled similarly to other insects. Stingless bee thorax mass scaled about isometrically (slope= 0.1.05 Adj. $R^2$=0.895, $\lambda$=0.858 [Table 1]), consistent with orchid bees (probably the most studied group of for flight physiology) and other bees and insects measured (Darveau et al., 2005; Dillon and Dudley, 2004). Abdomen mass also scaled isometrically (slope= 1.11 Adj. $R^2$= 0.86, $\lambda$=0.567[Table 1]) and head mass scaled hypometrically (slope= 0.86, Adj. $R^2$= 0.906, $\lambda$=0.00 [Table 1] ) as found for other insects and vertebrates (Darveau et al., 2005; Grula et al., 2021; Eberhard and Wcislo, 2011).

The differential scaling of flight costs in stingless and Euglossine bees was also supported by differential scaling of wing beat frequencies in these groups. In a strikingly different pattern than observed for fliers with larger masses, wing beat frequency was independent of mass (Fig. 2F, 3C). Most studies of insects have found that wing beat frequencies decrease with increasing body mass. For example, in Euglossines, the scaling exponent for wing beat frequency was -0.31 (Roberts et al., 1998). However, our results for a differential scaling of wing beat frequencies in smaller insects are supported by Byrne (1988), who demonstrated that aphids and white flies below 30 mg do not demonstrate reduced wingbeat frequencies in the larger species.

As for most other fliers studied (Lehmann, 2009, 1999; Marden, 1994), bees lifted similar fractions of their body mass during voluntary load-lifting of nectar, despite their varied thorax temperatures and hypermetric scaling of costs of flight when not loaded. Similarly, using a progressive load-lifting method, Dillon and Dudley found that vertical force production scaled either isometrically (using log-transformed data) or hypometrically (using raw data) across Euglossine bees (Dillon and Dudley, 2004). Thus, smaller stingless bees can carry similar loads (mass-specifically) at reduced cost relative to larger stingless bees.  Wing size scaled hypometrically (smaller bees had relatively larger wings) among stingless bee species (length width, and area; Fig 3 1, B) which likely contributes to the lack of scaling in wingbeat frequencies and load carriage.

Finally, we combined our data with all currently published data on flight metabolic rates of hovering insects to synthesize the scaling of flight costs across this clade (Table S1). The flight metabolic rates of stingless bees closely approximated costs of other similarly sized insects (Fig. 4). Inspection of all insect flight metabolic rates indicated that there was a breakpoint in the scaling of metabolic rates with size; a breakpoint analysis indicated that the breakpoint occurred at 58 mg (Fig. 4). A biphasic model using two size classes (above and below 58 mg) better explained the scaling of metabolic rates than a simple continuous log-log model, based on residual MSE of the generated breakpoint models compared to the standard model (Table 2). We next fit linear models to log-log plots of metabolic rates vs. mass above and below 58 mg; these had high $r^2$ values, particularly in the low range, (Table 2). The scaling slope of flight metabolic rate below 58 mg was 1.199, and 0.675 above 58 mg (Fig.4). We conclude that scaling of insect flight metabolic rates is biphasic, with hypermetric scaling in the low range and hypometric scaling in the high range.

The mechanisms responsible for the biphasic scaling of flight costs remain unclear, but likely include both aerodynamic and evolutionary mechanisms, probably working together.

Aerodynamic costs of flight may be reduced among smaller insects, partly due to performance at low Reynolds numbers. Smaller animals experience lower Reynolds number conditions greater air viscosity and less momentum, or inertial force). In these conditions, less energy would be required for flight simply because of the relatively increased viscosity as experienced by smaller flyers. The air would effectively increase the “buoyancy” of smaller animals much more than larger flyers. In these conditions, very tiny flying animals have evolved differences in flight strategy such as decreased wing veination or “feather” wings that allow them to float through air rather than actively flying (Farisenkov, 2020). Another possible strategy for flying in low Reynolds number conditions would be using wings like boat paddles. Smaller stingless bees may use their relatively larger wings to paddle more slowly than expected for their size through viscous air, which conserves energy. Additionally, Evolutionary mechanisms include the finding that smaller stingless bees have relatively larger wings (Fig. 3), as well as decreased veination on the laminar surface of the forewing, a relatively larger stigma, and a heavier forewing leading edge (Combes and Daniel, 2003; Danforth, 1989; Nel et al., 2012). These characteristics provide greater lift generation while gaining the energetic advantage of relatively low wing beat frequency as found here. Smaller stingless bees also have proportionally larger heads; this contributes to a shift in the center of mass to a more forward position (Ellington, 1984; Liu, 2009). Such morphological changes may contribute to using different aerodynamic mechanisms such as “clap and fling” found in some tiny species (Farisenkov et al., 2021, 2020; Miller and Peskin, 2009). Regardless of the mechanism, the reduced cost of flight at sizes below 50 mg will likely reduce costs of flight for foraging, defense, and migration, providing a significant selective advantage for the evolution of small body size among insects. However, it should be noted that these reduced costs for small insect fliers applies to hovering flight as studied here, when costs of generating lift predominate. During forward flight, drag forces on the body become important, and the low Reynold's number and high surface-to-volume ratios of smaller insects may cause an increase in flight costs (Ellington, 1999, 1985; Fry et al., 2005).

The biphasic scaling of flight metabolic rates in insects seems to contradict predictions of hypotheses for metabolic scaling that depend on supply constraints in larger animals. Also, this biphasic pattern further reinforces the growing consensus that scaling patterns are variable with clade and type of activity. At least for insect hovering flight at body masses from 1 – 1000 mg, our data support the hypothesis that patterns of metabolic scaling are determined by changes in the aerodynamic cost of locomotion, and changes in skeletal muscle cycle frequency.

**Acknowledgments**

We acknowledge Panama's Ministerio de Ambiente and the Smithsonian Tropical Research Institute (STRI) for permitting and aiding in this research. We are especially grateful to the following individuals at STRI for logistical and technical support: Bill Wcislo, Jon Shik, Lil Camacho, Raineldo Urriola. Paola Galgani, Vilma Fernandez, and Oris Acevedo. We are also grateful to Josh Borup who contributed to wing morphology data collection.

**Funding**

This work was supported by the Society of Integrative & Comparative Biology (SICB), the National Science Foundation (IOS 1122157 to J.F.H), and the Smithsonian Tropical Research Institute.

### Figures

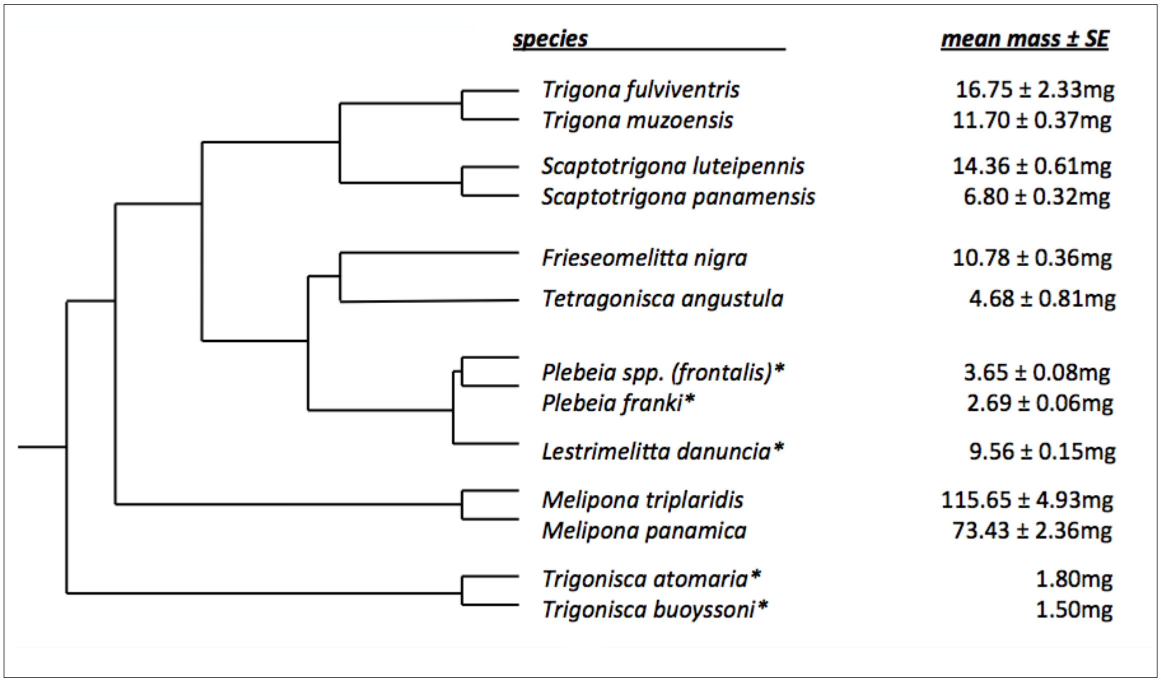


**Figure 1.** Phylogenetic tree of stingless bee species included in this study based on relationships found in Rasmussen & Cameron 2007 and 2009. All branch lengths are set equal to one because of the absence of some species from available molecular phylogenies of Meliponines. Phylogenetic independent contrasts demonstrate that phylogeny is not a significant factor in our analysis (Table 1). *Plebeia spp. (frontalis)* is undescribed. PGLS analysis was done with and without this species included and did not yield different results. Average body mass ±SE are indicated next to species names and miniaturized lineages are specified with an asterisk according to Michener (2001) and Camargo (2013).

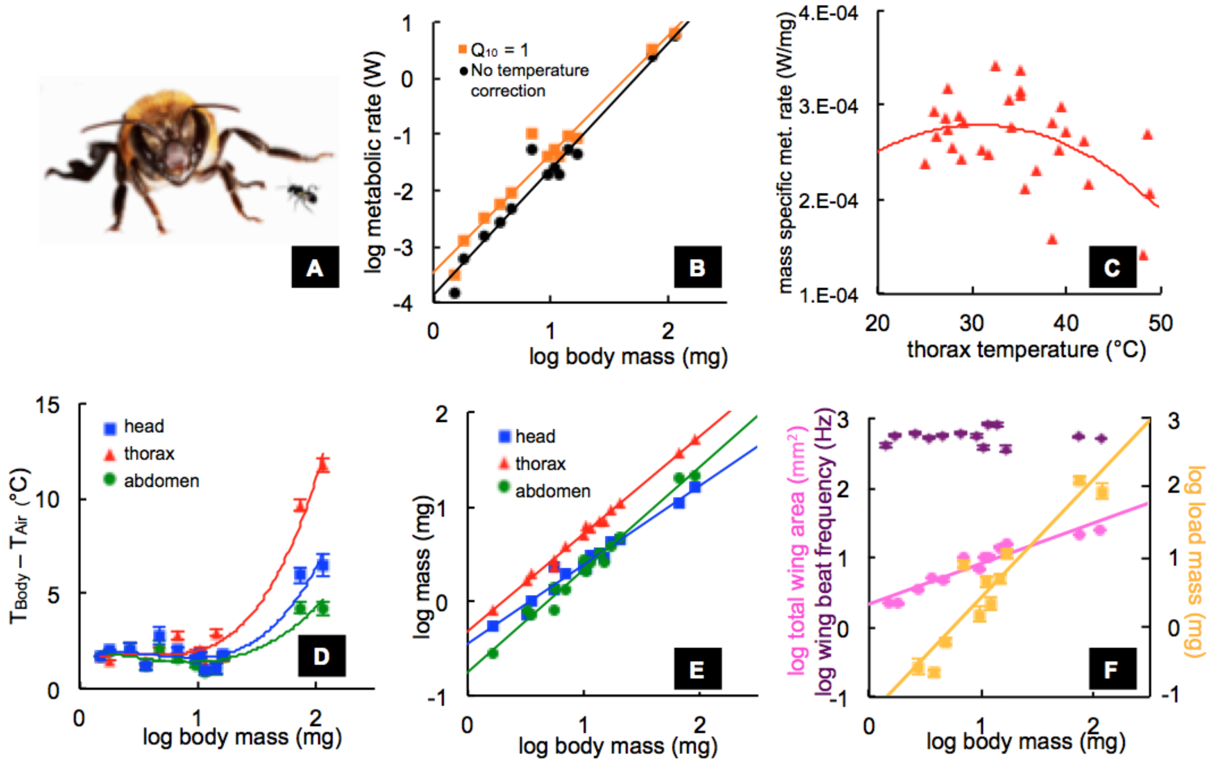


**Fig. 2 (A)** Size comparison of biggest and smallest stingless bees included **(B)** Metabolic rates of stingless bees with and without $Q_{10}$ correction of 2. Line shows second order polynomial fit. **(C)** The thermal flight performance curve of *S. luteipennis* (n=30) indicates that $Q_{10}$ for this stingless bee is approximately equal to 1. **(D)** Body segment temperature elevation above air temperature. Small bees (< 20 mg) had body temperatures 0.7-3°C above air temperatures, while large species (> 70 mg) had substantially elevated body segment temperatures. Lines show third order polynomial fits. **(E)** Thorax, head, abdomen mass scaled hypometrically (slopes= 1.05 for thorax, slope= 0.86 for head and 1.11 for abdomen) with body mass across stingless bees. **(F)** Wingbeat frequency was constant across body size while load carriage abilities scaled isometrically (slope =1.05). Total wing area scaled hypometrically, indicating that smaller stingless bees have proportionally larger wings. All multi species regression lines were plotted with PGLS.

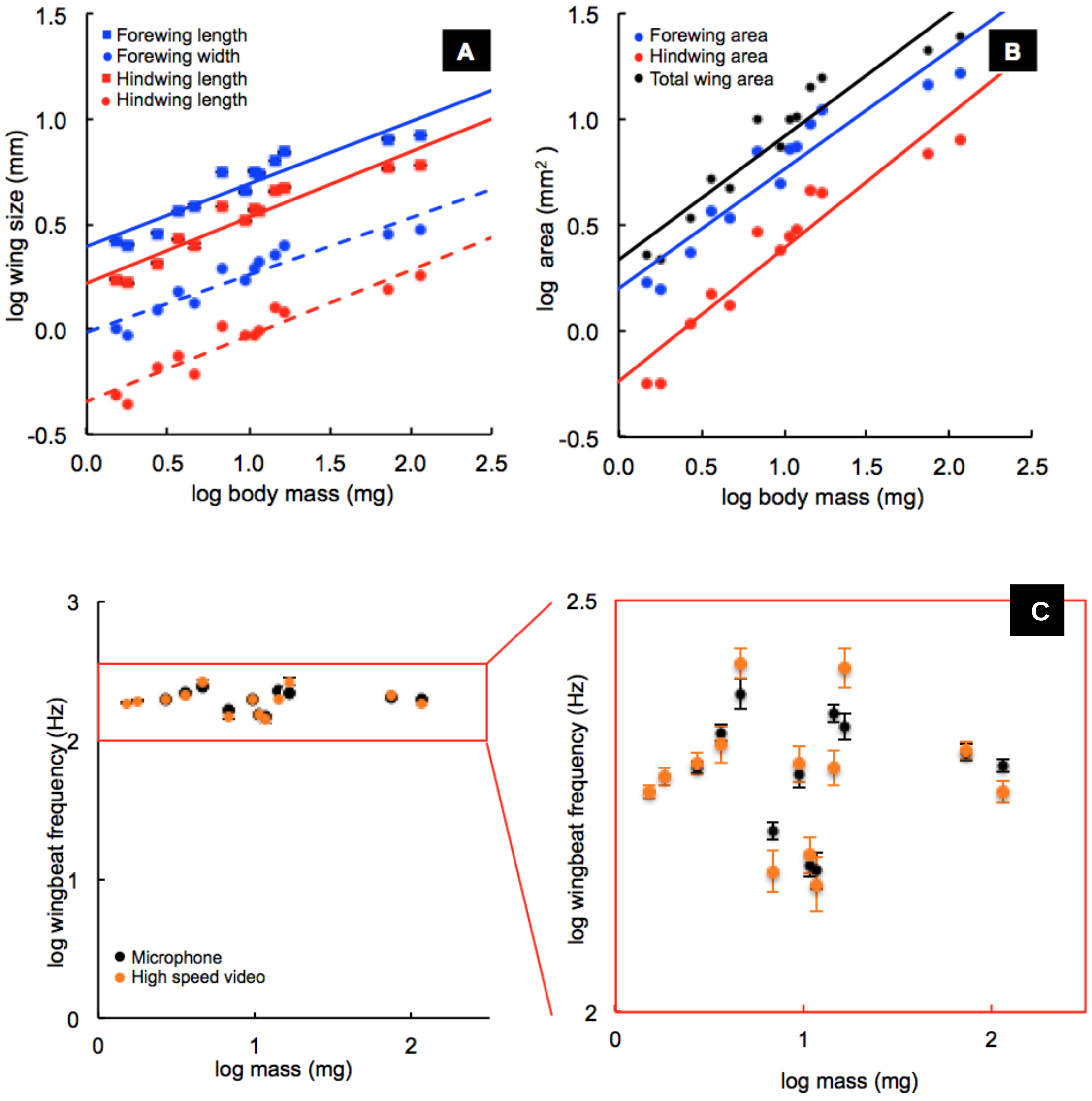


**Figure 3. (A)** Scaling relationships of forewing (blue) and hindwing (red) length (squares), and widths (circles). All scaled hypometrically with body mass (Table S1). **(B)** Scaling of total wing area (black), forewing area (blue), and hindwing area (red) with body mass. Forewings are proportionally larger in smaller bees than hindwings. All scaling parameters are listed in Table S1. **(C)** Comparison of microphone and high-speed video methods of wing beat frequency measurement. There was no significant relationship between mass and wing beat frequency among all species (slope= 0.02, Adj. $R^2$=0.066, $P$=0.524). Average wing beat frequency across species = 204.6 ± 8.3 SE beats/sec. Each point represents the average wing beat frequency within a species ±SE.

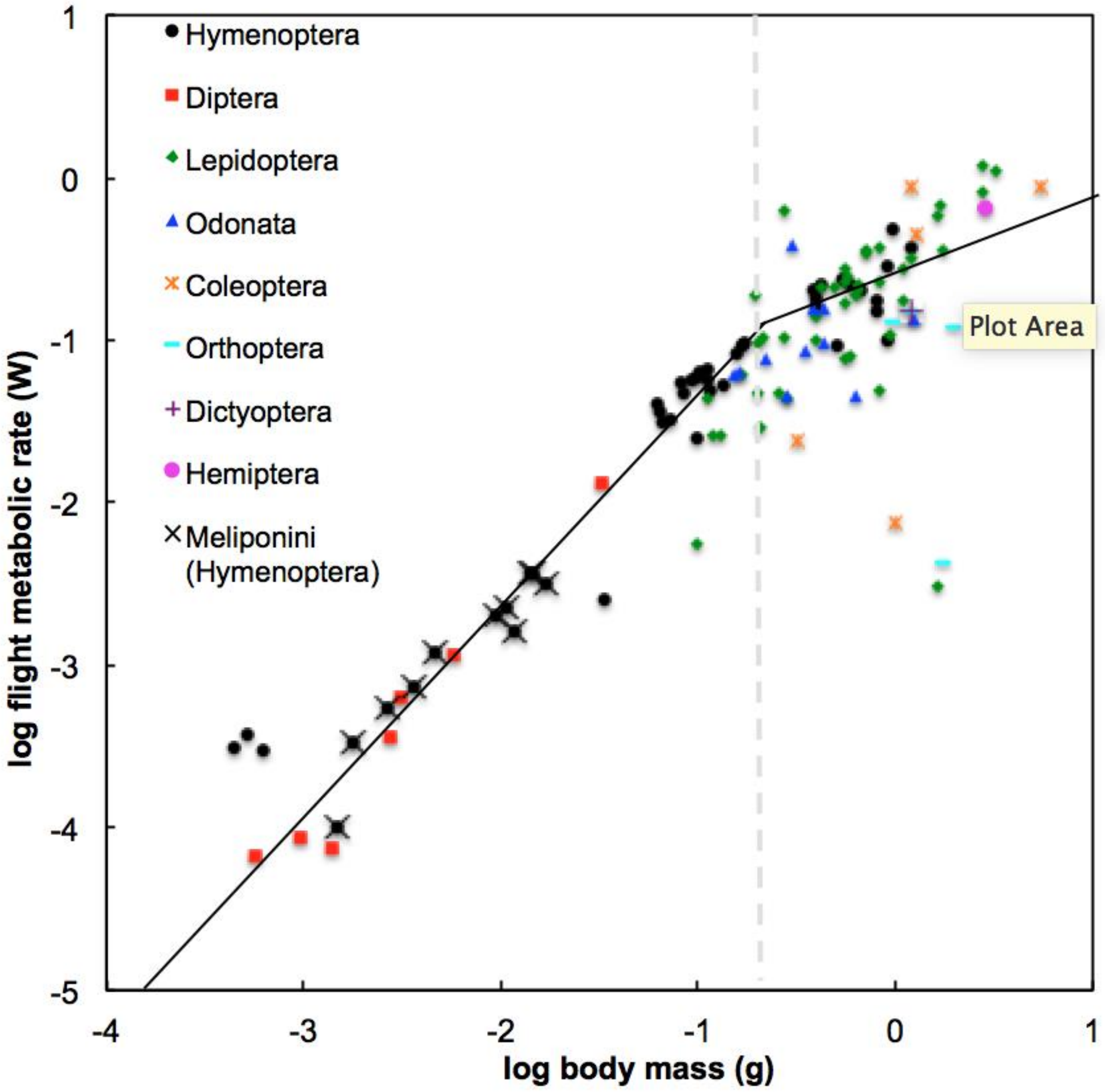


**Fig. 4.** Flying while small costs less. Flight metabolic rate in insects below 58 mg in body mass scales hypermetrically (slope =1.19) while flight metabolic rate in insect greater than 58 mg scales hypometrically (slope = 0.67) as traditionally observed among most animals.

**Tables**

**Table 1.** Phylogenetic generalized least squares regression (PGLS) statistics for all physiological variables for 13 species of stingless bees. All scaling data were regressed using PGLS as part of the regression model, which integrates linear models to fit a line based on evolutionary relatedness through data. Phylogenetic signal (λ) is on a 0-1 scale where 1 is the highest amount of signal possible. Coefficient t measures the distance of the line estimate (slope) from zero, with higher number demonstrating higher significance of the relationship between variables (body mass and the physiological variables shown). Slopes indicated in bold are not isometric. Other bolded values are statistically significant. Confidence limits are shown with 12 df (13 species included in analysis)

| measurement | slope | intercept | t | p | adj. $r^2$ | λ | Conf. Limits |
|---|---|---|---|---|---|---|---|
| *flight MR ($CO_2$ ml/h)* | **2.234** | 2.843 | 14.466 | **1.668e-08*** | 0.946 | 0.000 | 0.274 |
| *head temp(°C)* | 0.225 | 0.804 | 1.828 | 0.095 | 0.163 | 0.490 | 0.219 |
| *thorax temp (°C)* | 0.298 | 1.043 | 2.466 | **0.032*** | 0.297 | 1.000 | 0.216 |
| *abdomen temp (°C)* | 0.146 | 0.578 | 1.322 | 0.213 | 0.059 | 0.570 | 0.198 |
| *wing beat freq (Hz)* | 0.015 | 2.323 | 0.430 | 0.675 | -0.073 | 0.000 | 0.062 |
| *load carriage (mg)* | 0.869 | 0.776 | 15.666 | **7.693e-08*** | 0.967 | 0.000 | 0.100 |
| *total wing area ($mm^2$)* | **0.499** | 0.431 | 8.749 | **2.762e-06*** | 0.863 | 0.898 | 0.119 |
| *forewing area ($mm^2$)* | **0.567** | 1.869 | 8.308 | **4.55e-06*** | 0.850 | 0.832 | 0.121 |
| *hindwing area ($mm^2$)* | **0.643** | 1.639 | 9.757 | **9.449e-07*** | 0.887 | 1.000 | 0.118 |
| *forewing width (mm)* | 0.269 | 0.783 | 8.234 | **4.963e-06*** | 0.848 | 0.860 | 0.059 |
| *forewing length (mm)* | 0.301 | 1.281 | 7.767 | **8.646e-06*** | 0.832 | 0.842 | 0.069 |
| *hindwing length (mm)* | 0.314 | 1.147 | 8.459 | **3.826e-06*** | 0.855 | 0.775 | 0.066 |
| *hindwing width (mm)* | 0.315 | 0.585 | 8.110 | **5.733e-06*** | 0.844 | 0.904 | 0.069 |
| *head mass (mg)* | **0.860** | -0.921 | 9.839 | **4.097e-06*** | 0.906 | 0.000 | 0.155 |
| *thorax mass (mg)* | 1.046 | -0.235 | 9.275 | 6.668e-06 | 0.895 | 0.858 | 0.201 |

**Table 2.** Comparison of most likely linear and breakpoint models for flight metabolic rate scaling across flying insects. The breakpoint model has much higher support using Akaike's information criterion (AIC) than the standard linear model.

| Model | Slope | St. Err. | p (slope = 0) | p (slope = 0.67) | p (slope = 1) | Intercept | θ | AIC | Akaike weight |
|---|---|---|---|---|---|---|---|---|---|
| **Linear** | 0.98 | 0.04 | **<0.001*** | NA | 0.562 | -0.62 | NA | 139.03 | 0.003 |
| **Break-point** | Left: 1.19<br>Right: 0.67 | 0.07<br>0.17 | **<0.001***<br>**0.004*** | NA<br>0.305 | **0.027***<br>**0.004*** | -1.03 | -0.63 | 127.34 | 0.997 |
