## Supplemental figure 1 for "Isometric Spiracular Scaling in Scarab Beetles: Implications for Diffusive and Advective Oxygen Transport"

### Supplementary Figure 1

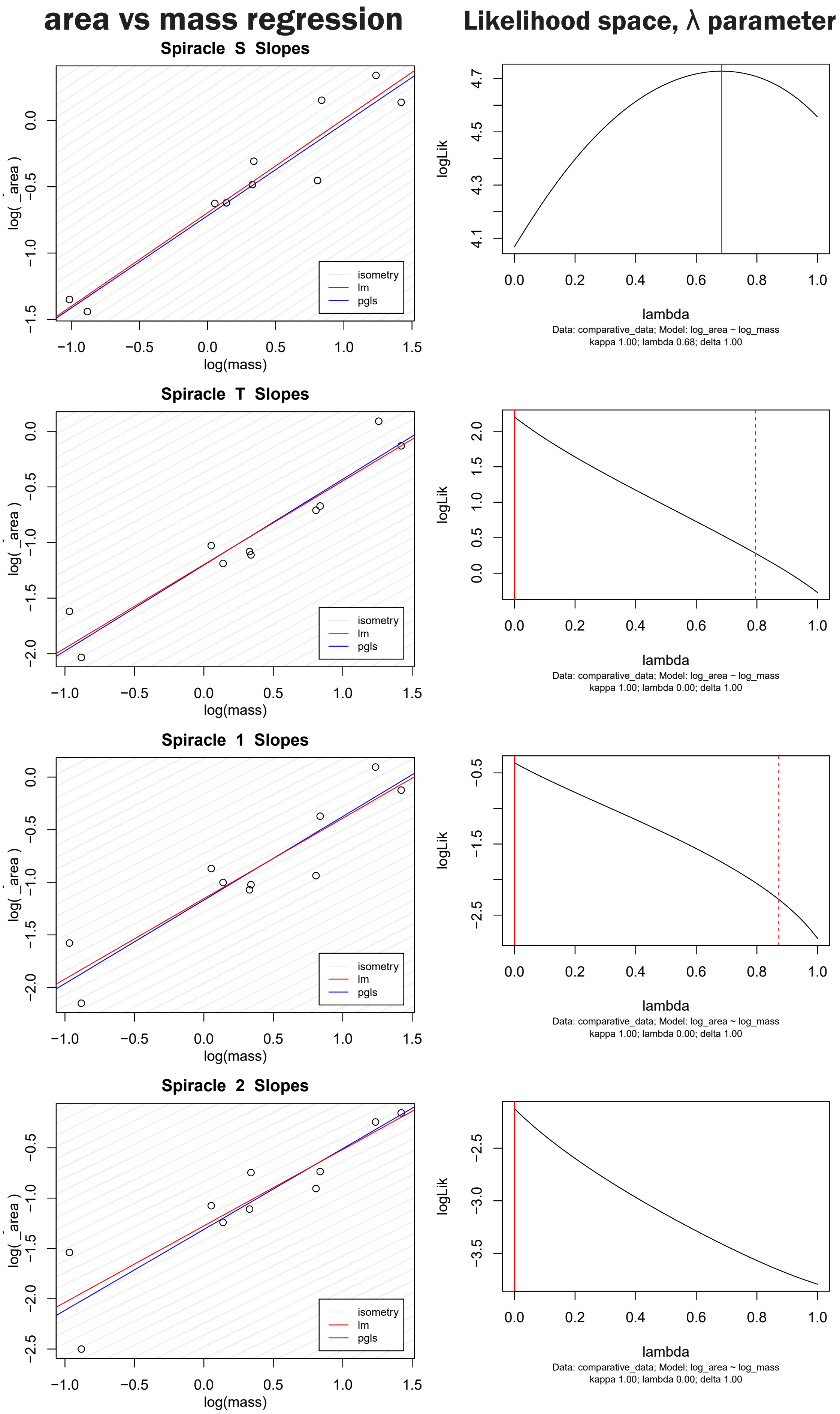

### area vs mass regression

### Likelihood space, λ parameter

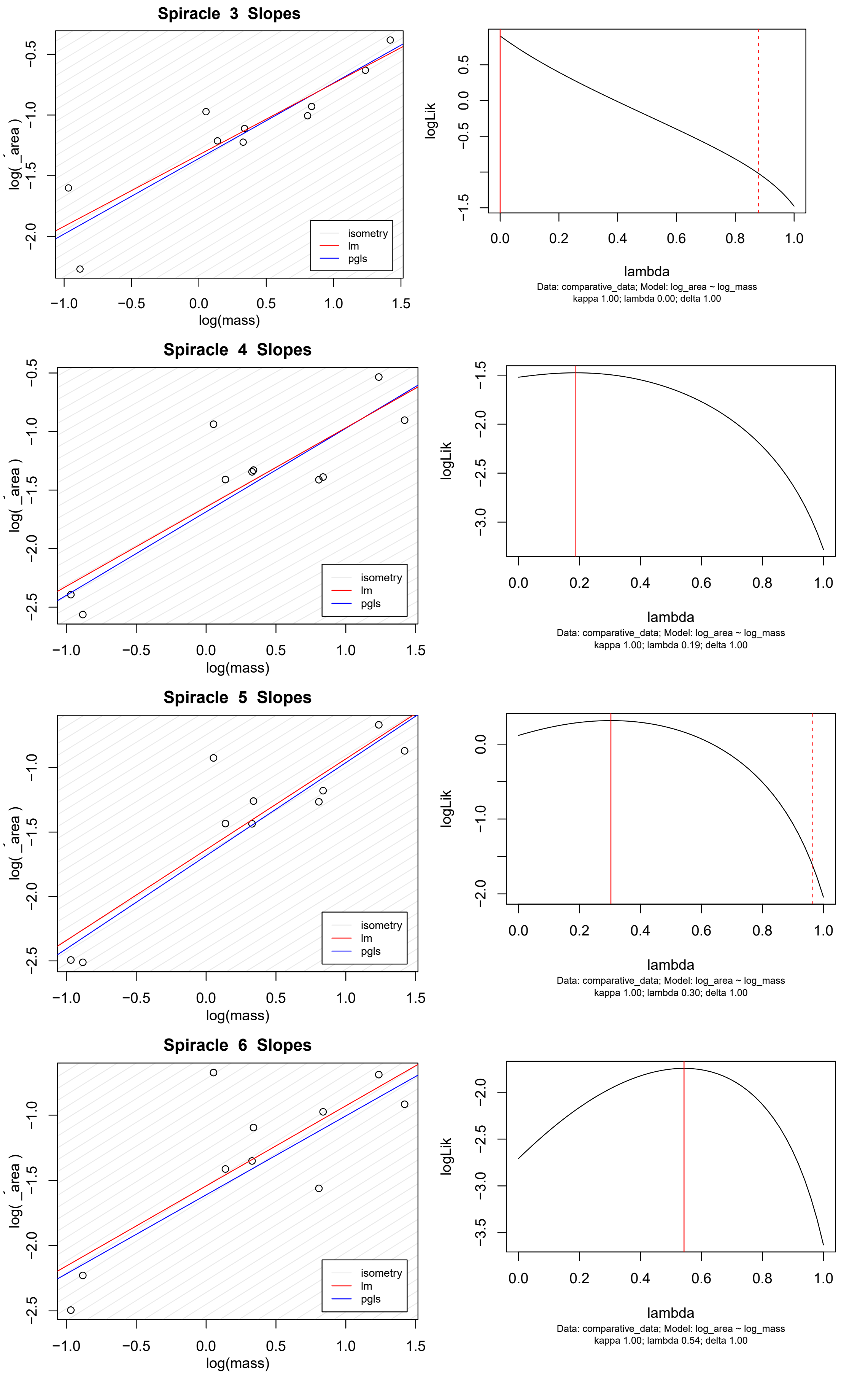

### depth vs mass regression

### Likelihood space, λ parameter

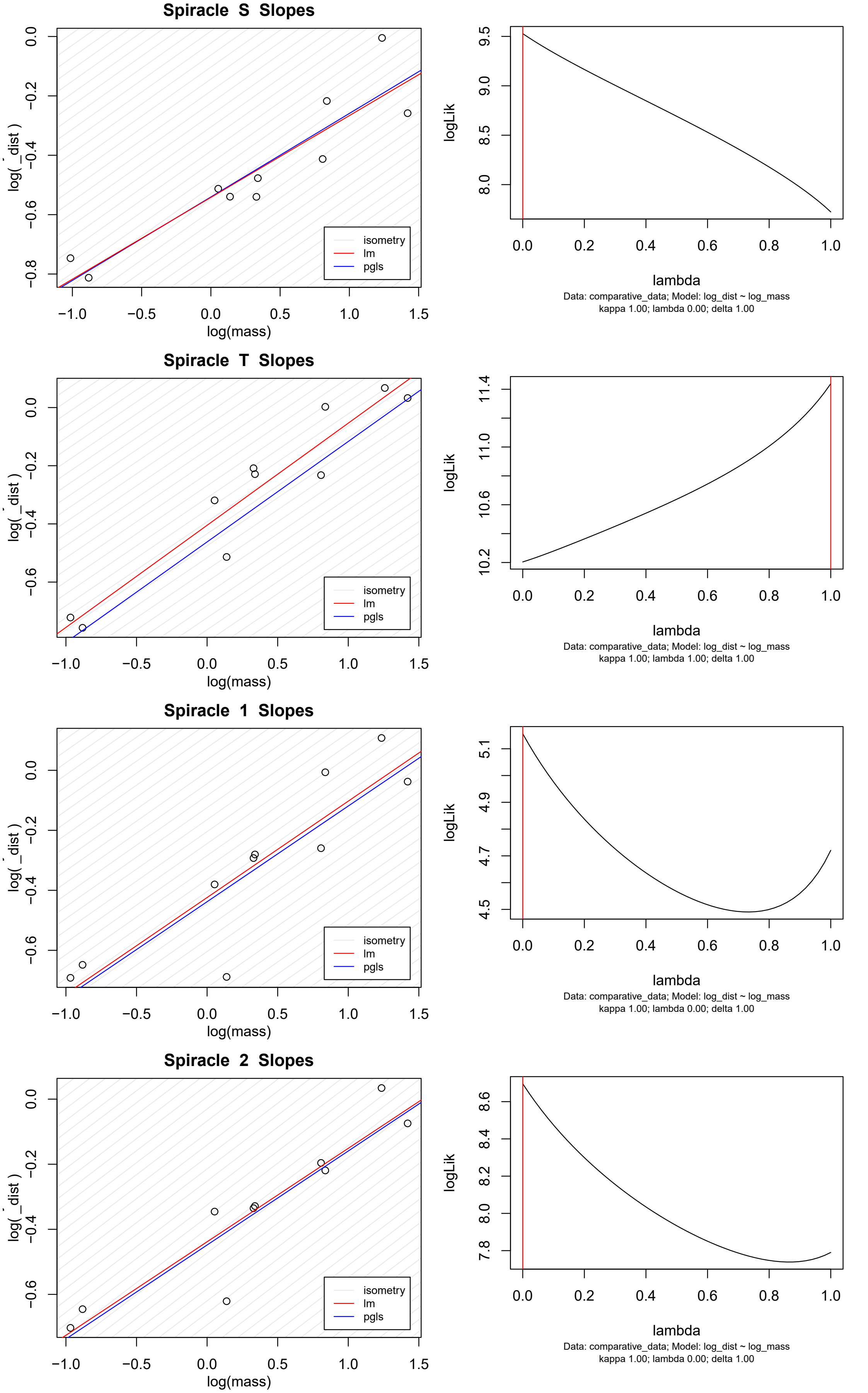

#### depth vs mass regression

#### Likelihood space, λ parameter

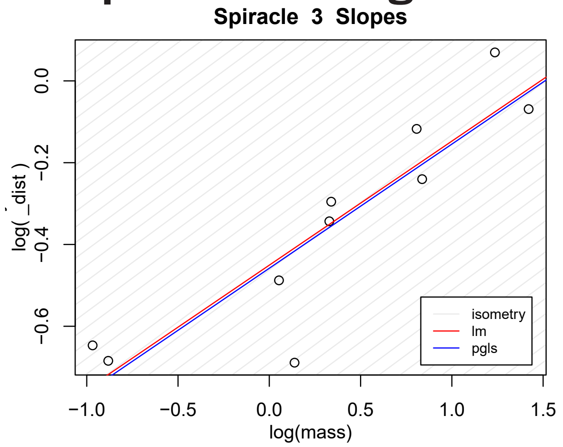


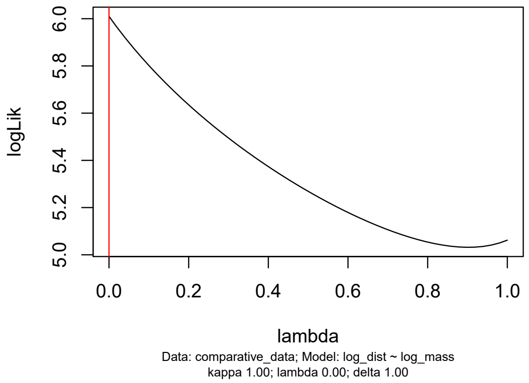


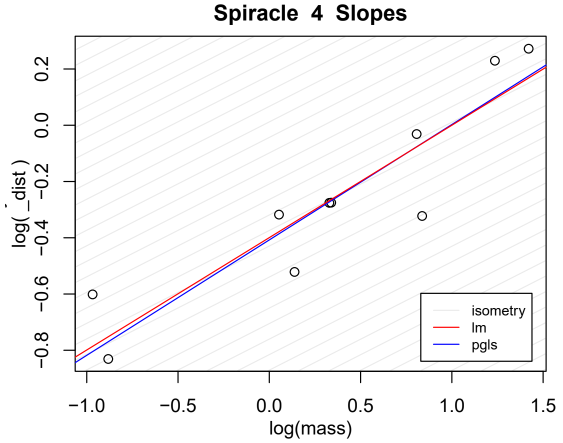


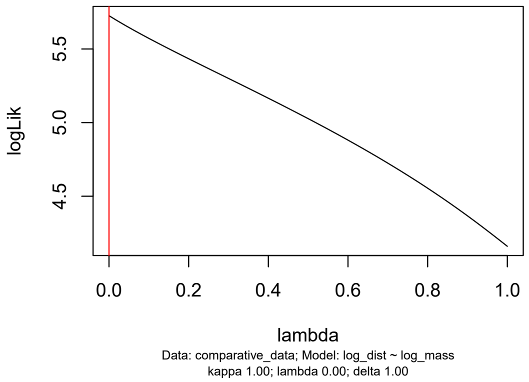


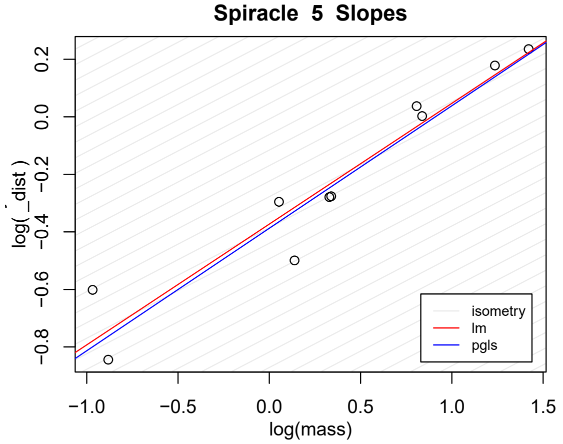


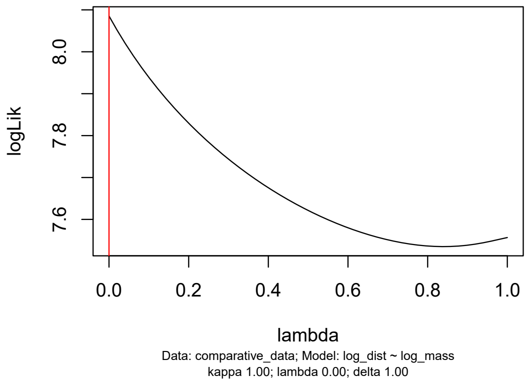


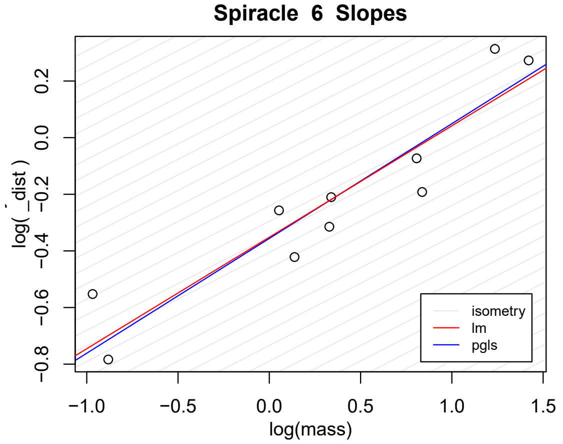


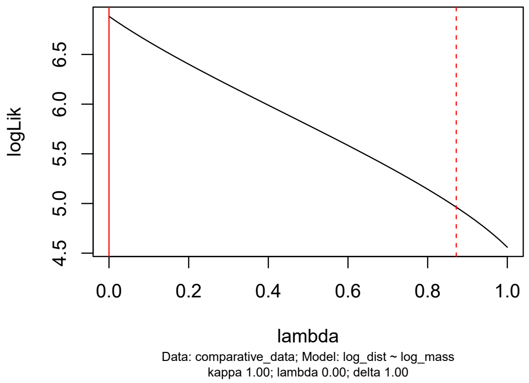

### area/depth vs mass regression

### Likelihood space, λ parameter

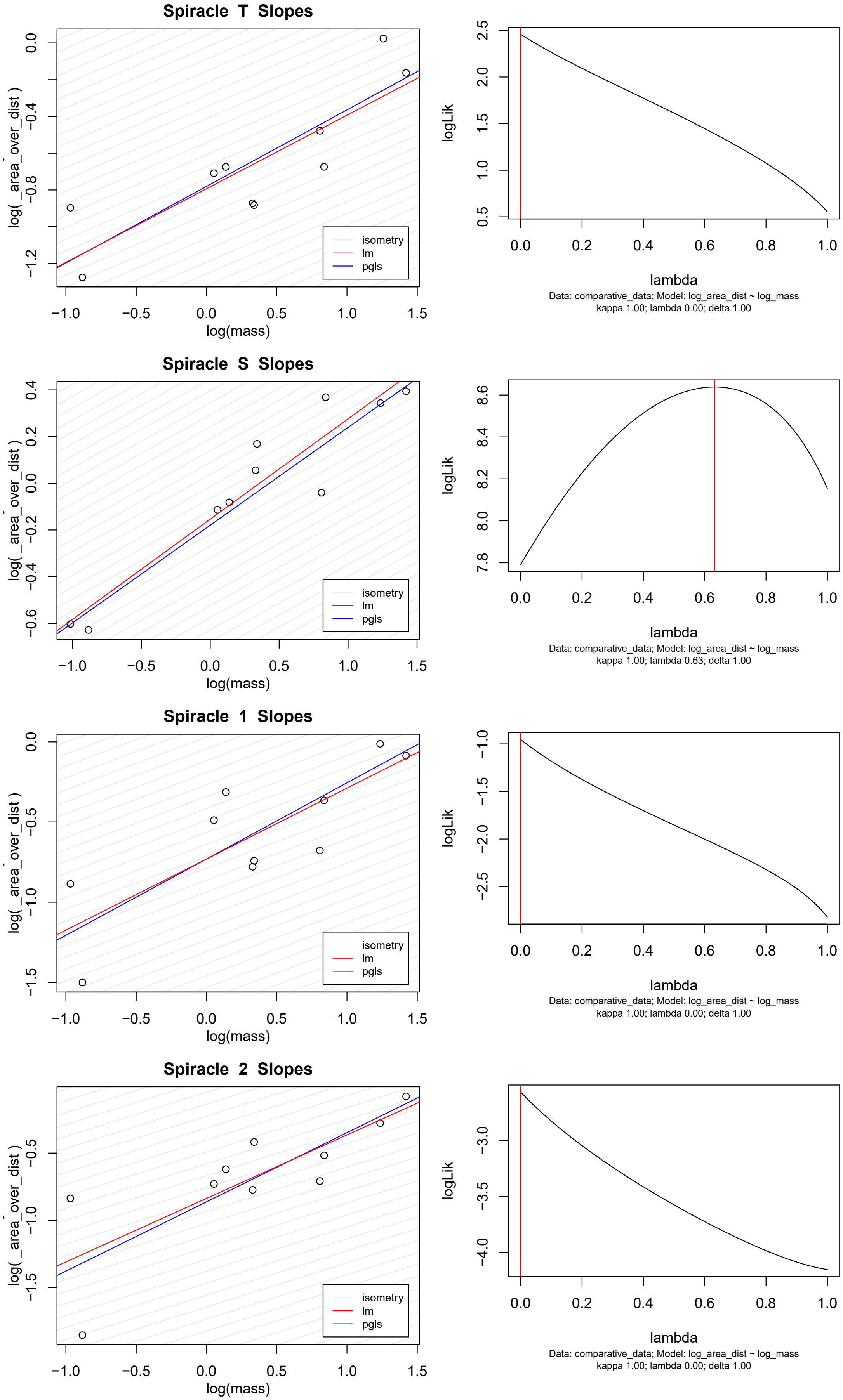

### area/depth vs mass regression

### Likelihood space, λ parameter

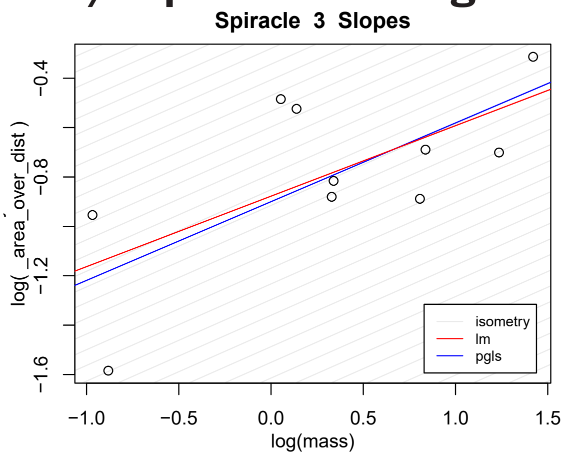


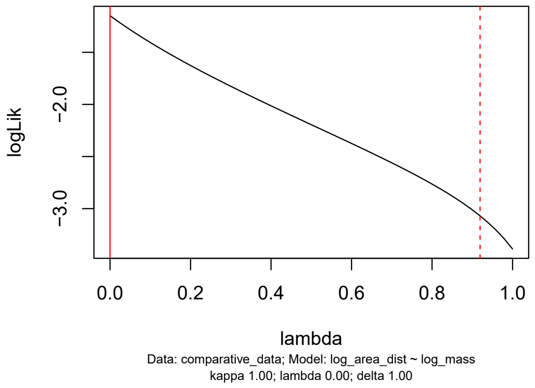


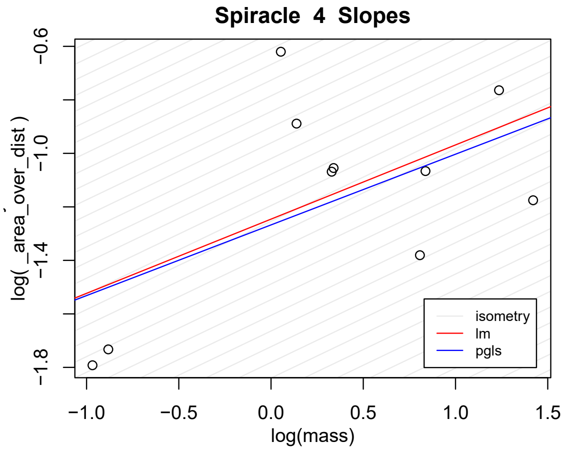


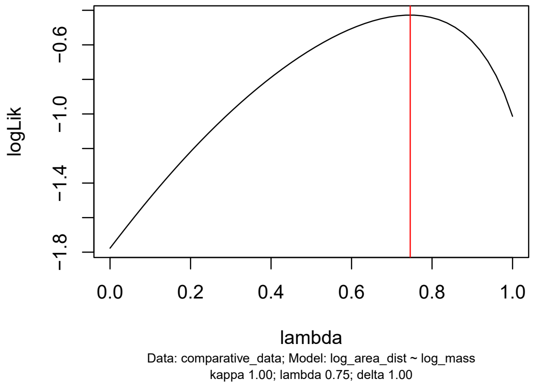


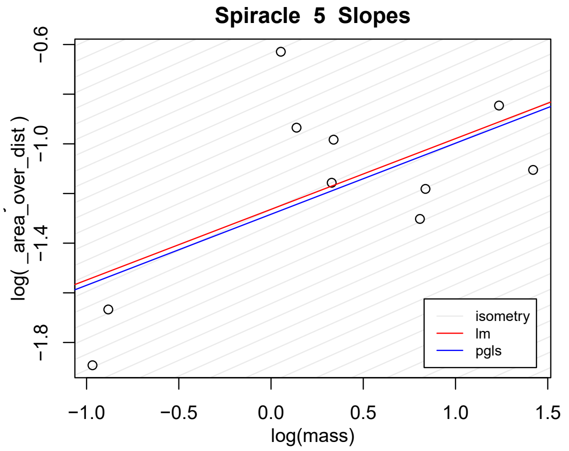


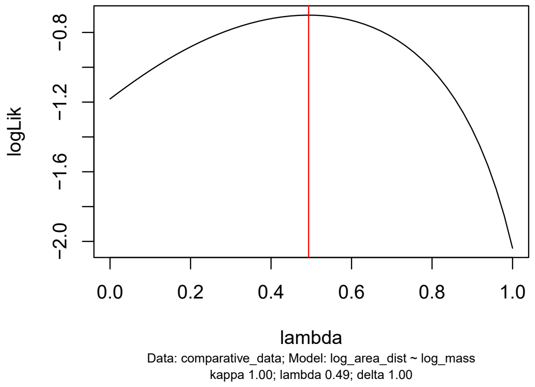


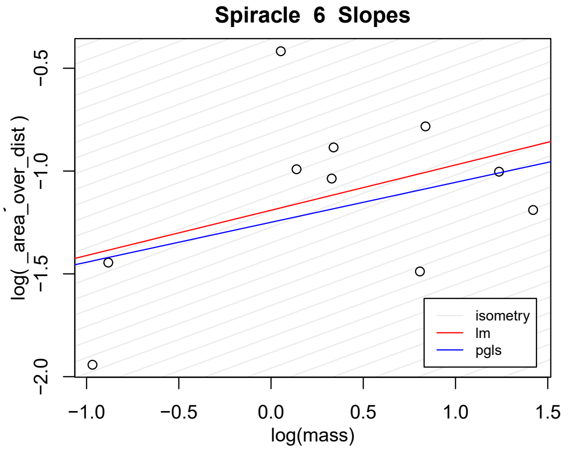


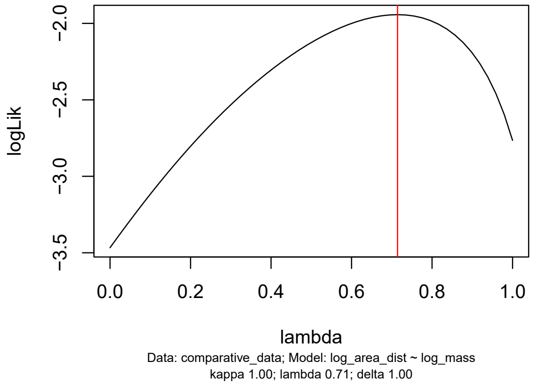

#### area$^2$/depth vs mass regression | Likelihood space, λ parameter

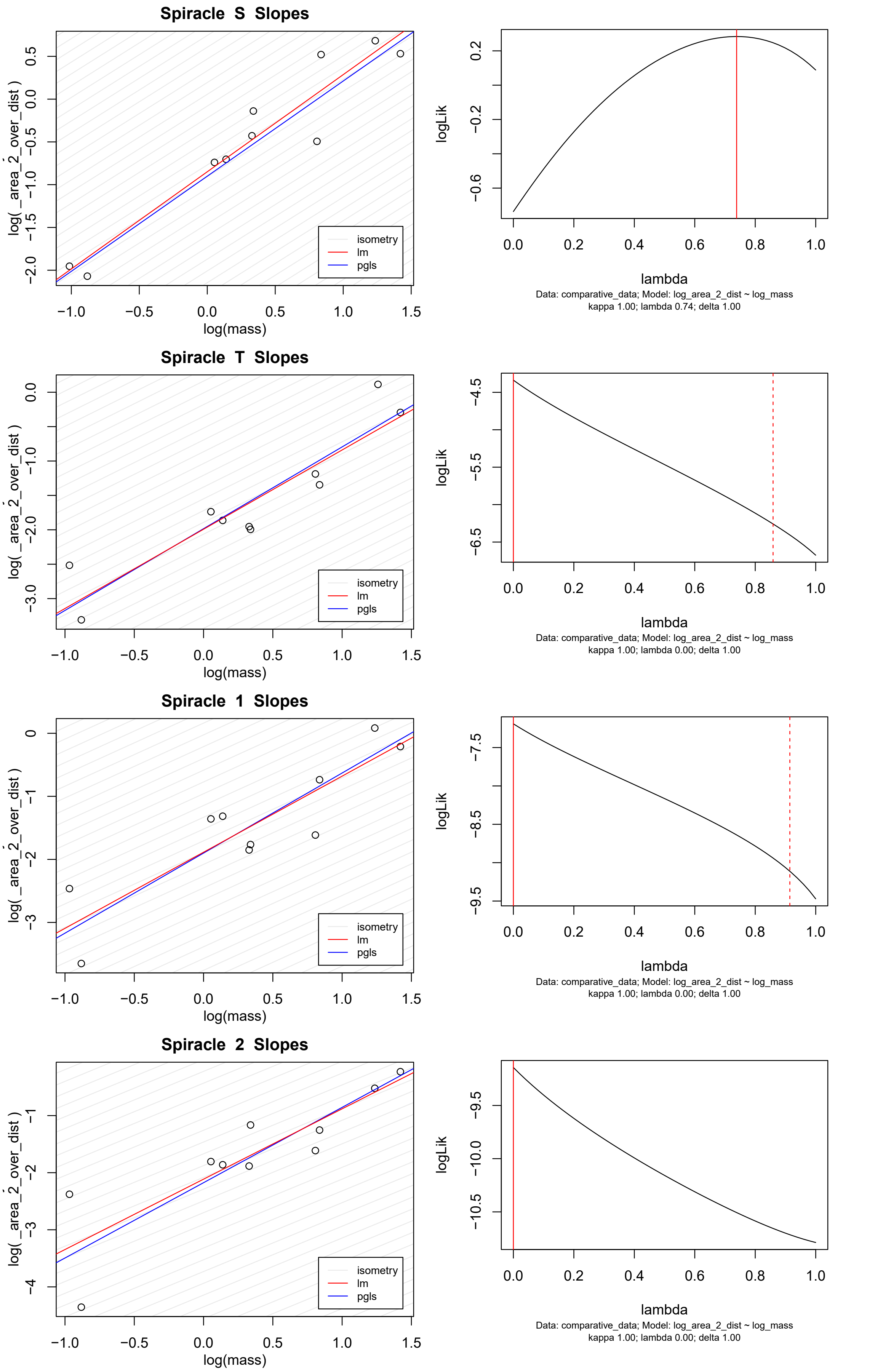

### area²/depth vs mass regression

### Likelihood space, λ parameter

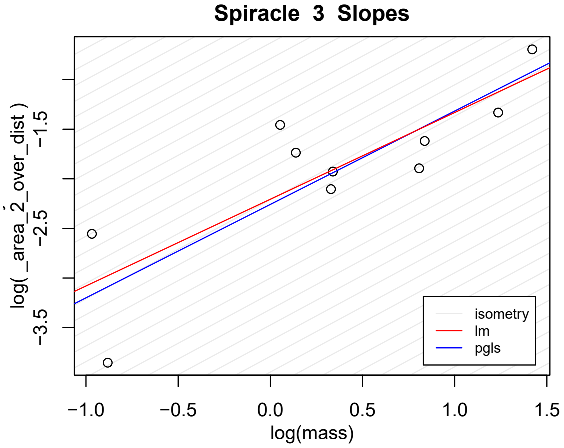


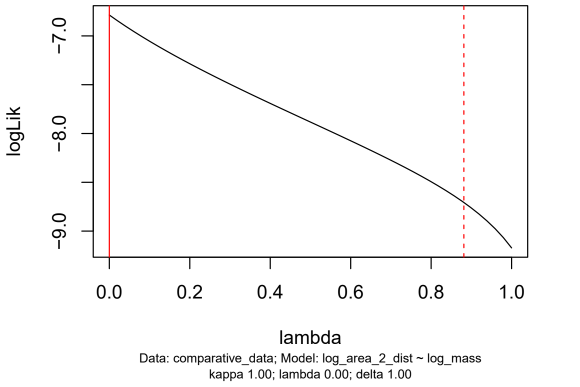


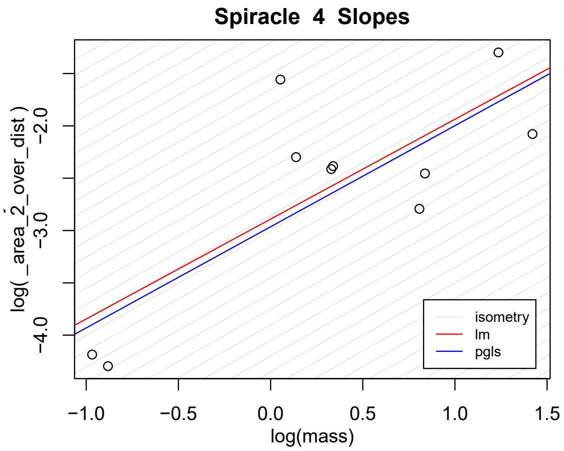


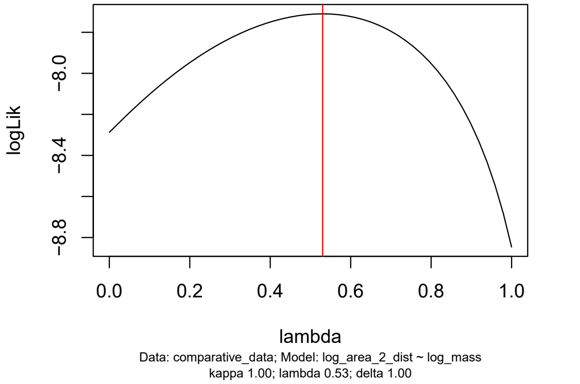


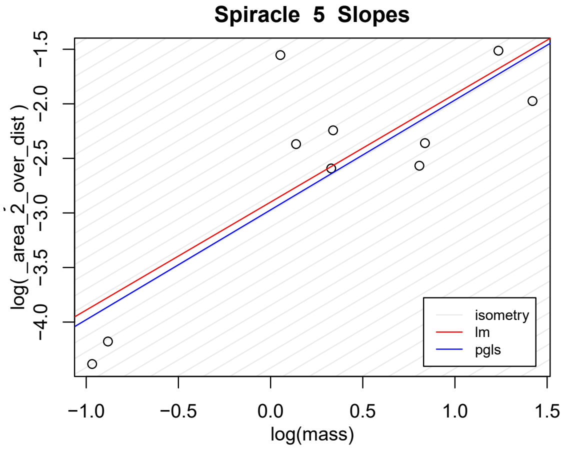


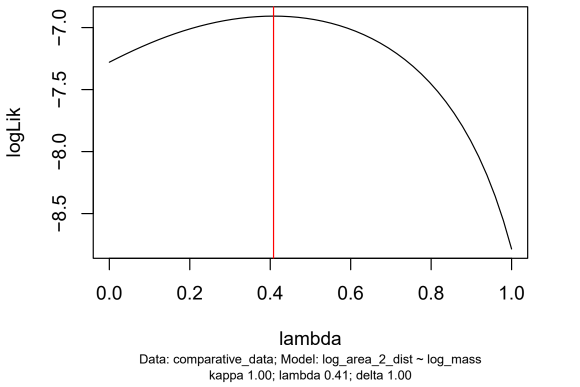


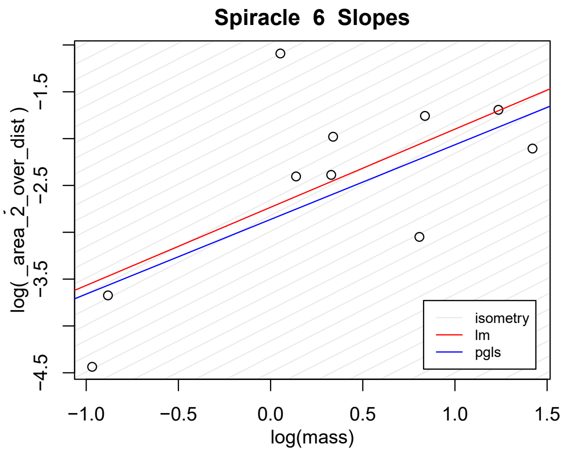


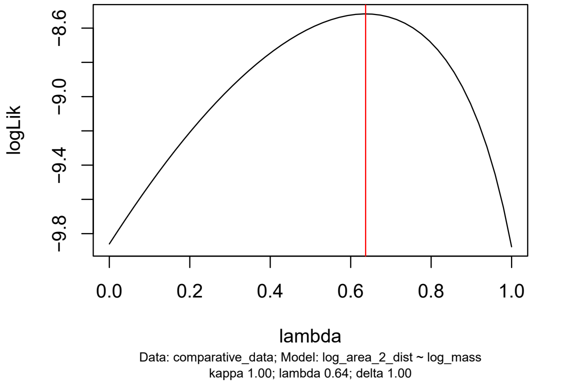
