## supplemental figure 2 for "Isometric Spiracular Scaling in Scarab Beetles: Implications for Diffusive and Advective Oxygen Transport"

Supplementary Figure 2

### Nonidentifiability of phylogenetic signal parameter, area vs mass

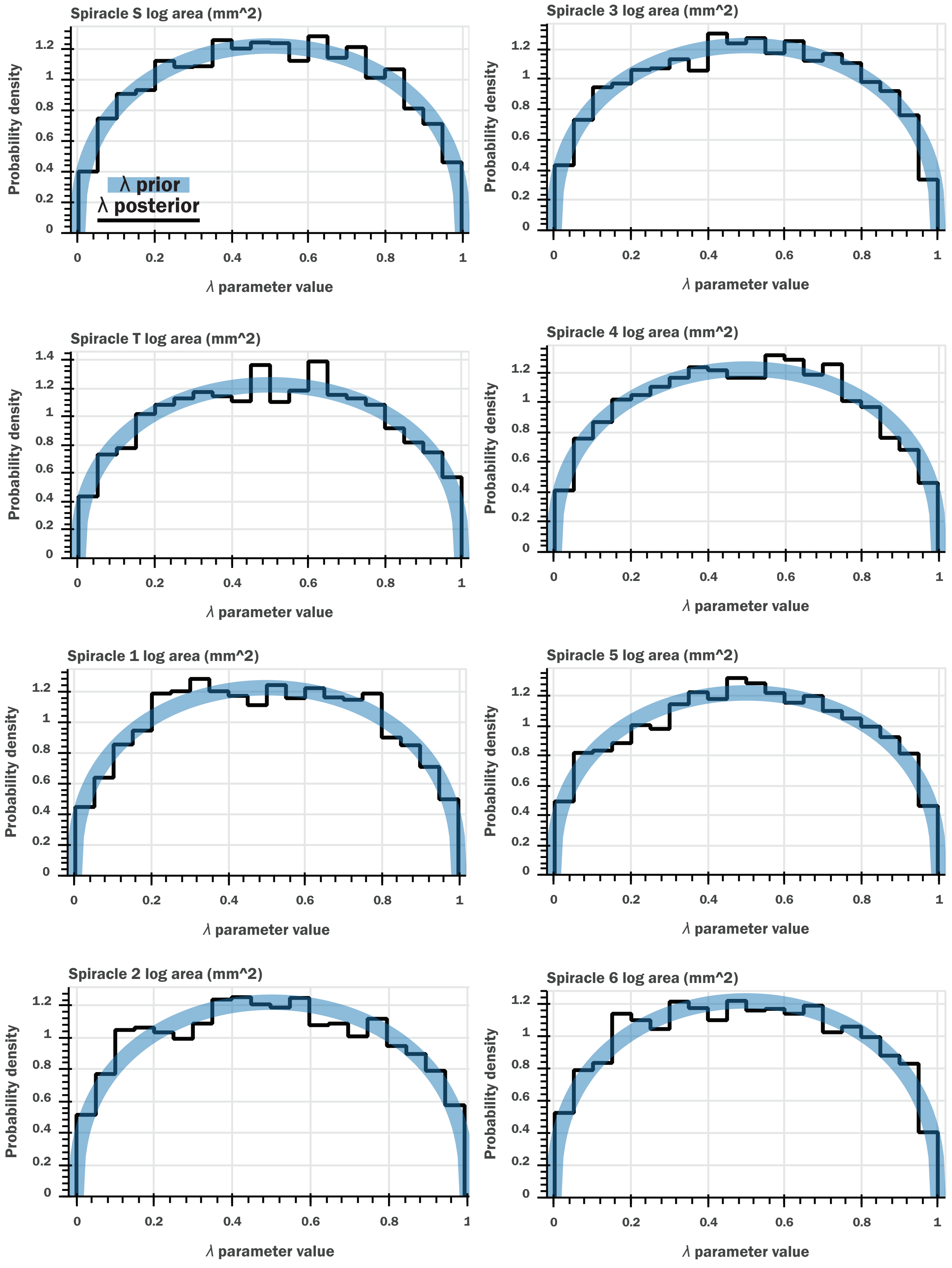

### Nonidentifiability of phylogenetic signal parameter, depth vs mass

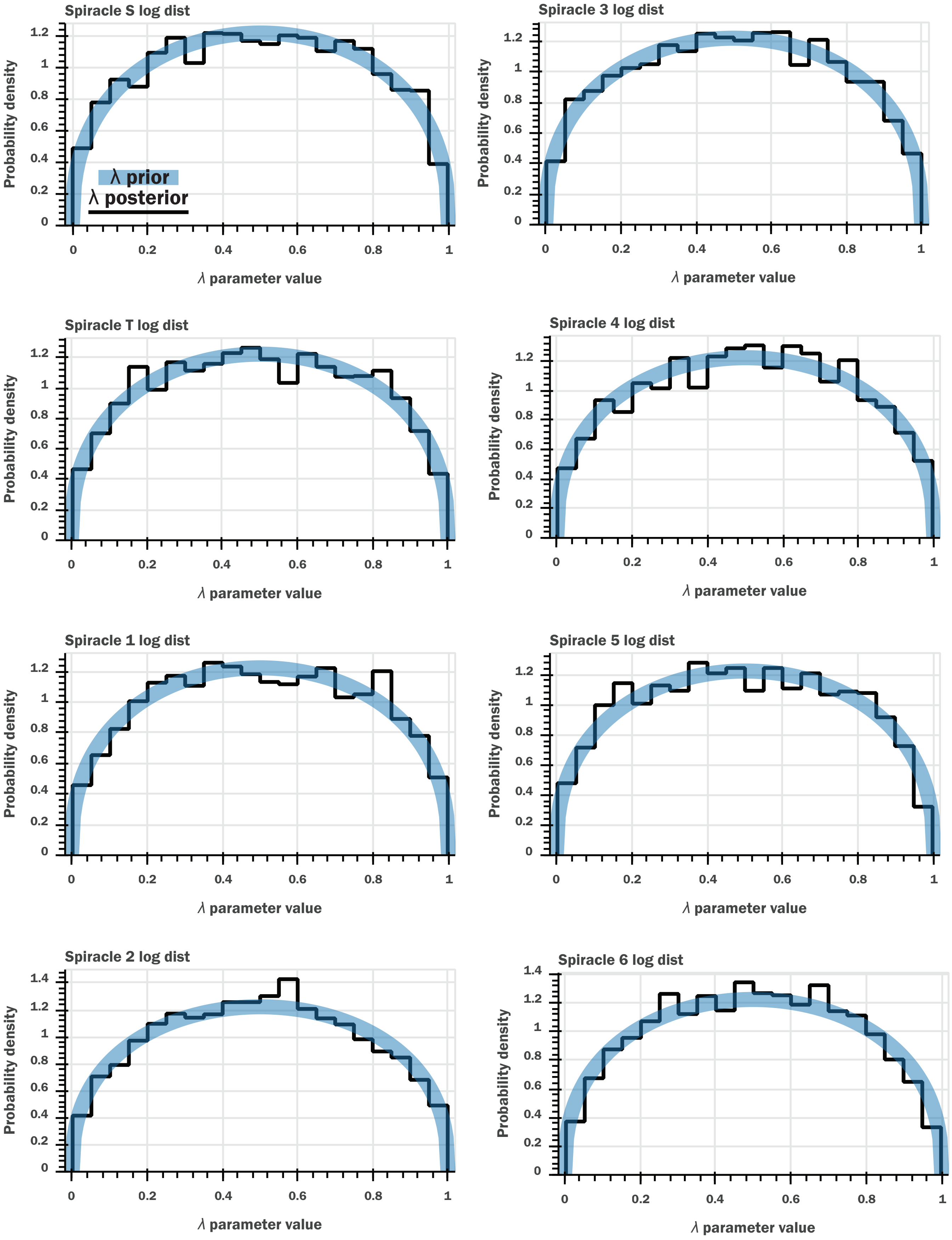

### Nonidentifiability of phylogenetic signal parameter, area/depth vs mass

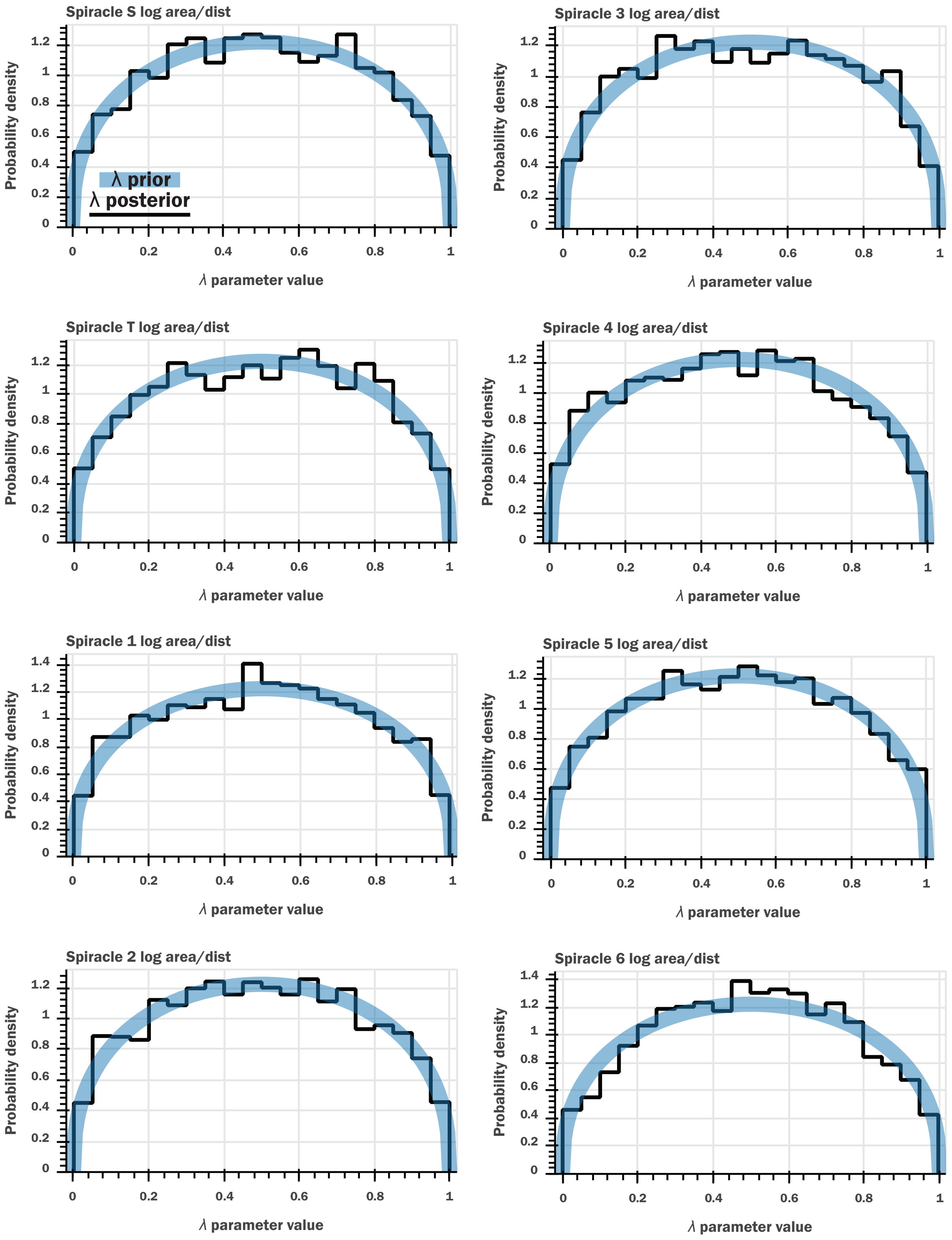

### Nonidentifiability of phylogenetic signal parameter, area²/depth vs mass

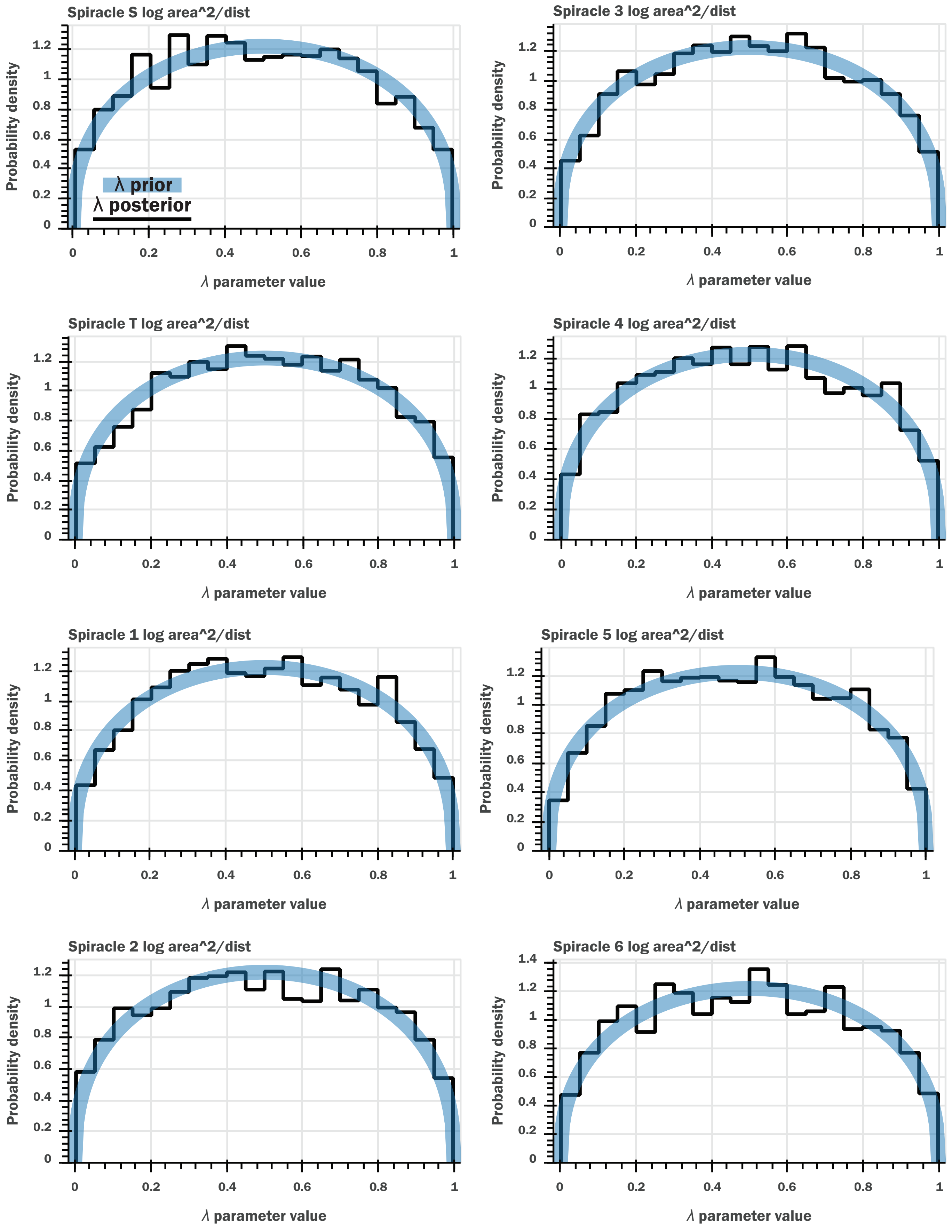
