## Supplementary figures and images for "Isometric Spiracular Scaling in Scarab Beetles: Implications for Diffusive and Advective Oxygen Transport"

### supplemental figure 3

**Supplementary Figure 3**

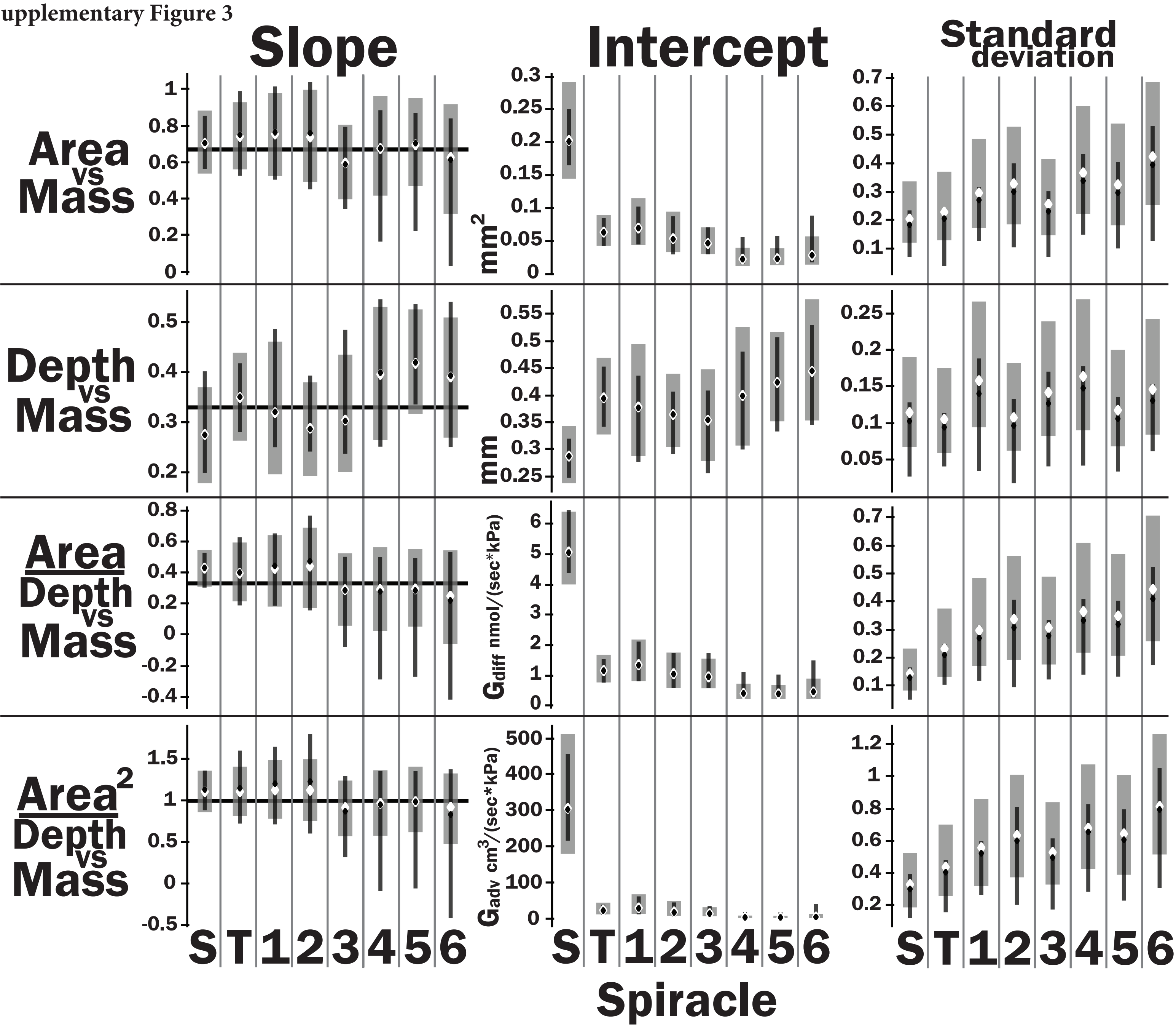
