## supplemental figure 4 for "Isometric Spiracular Scaling in Scarab Beetles: Implications for Diffusive and Advective Oxygen Transport"

Supplementary Figure 4

### Bayesian regression, area vs mass

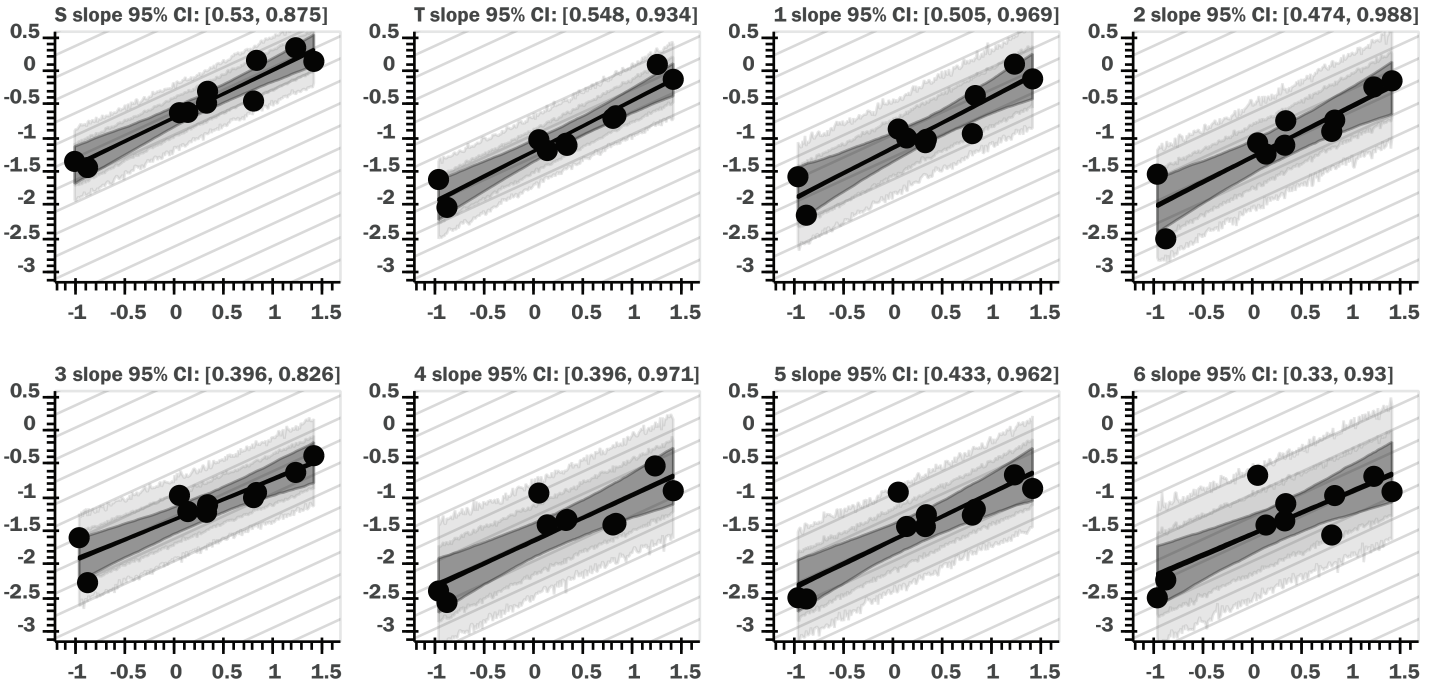


### OLS regression, bootstrap CI, area vs mass

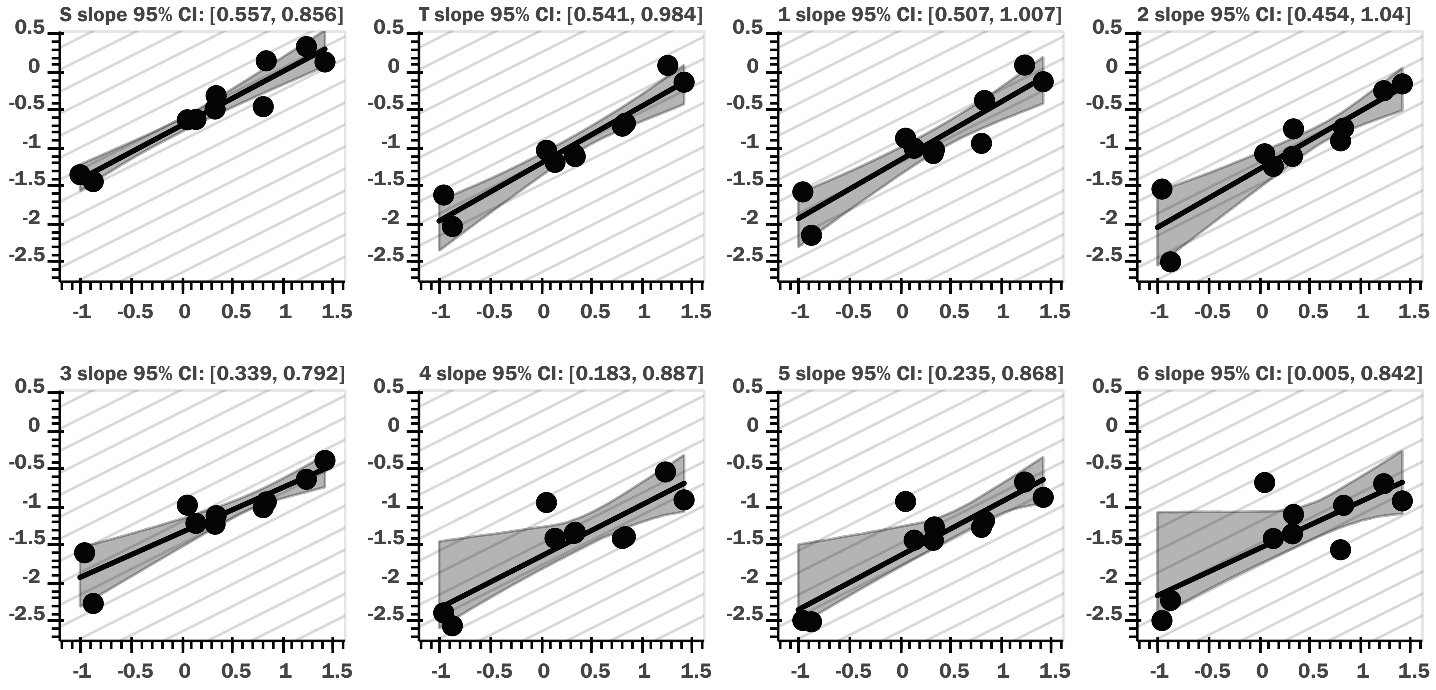

### Bayesian regression, depth vs mass

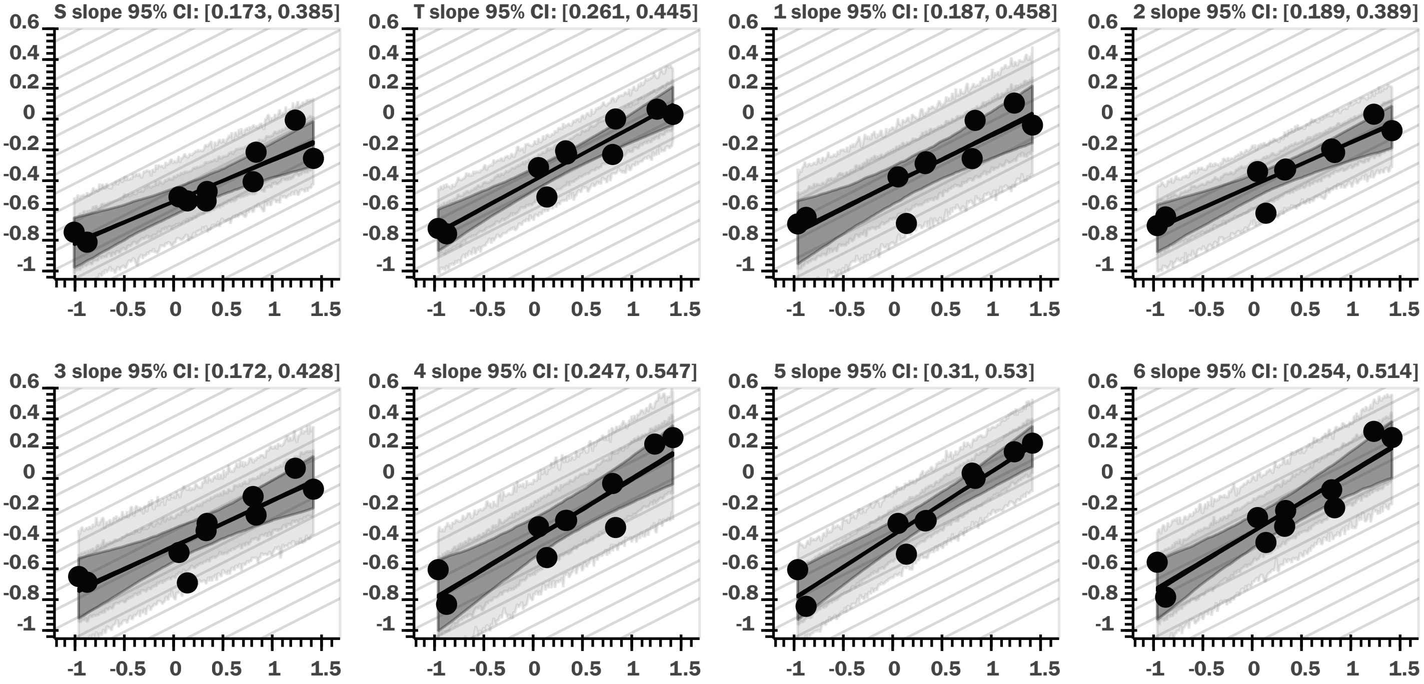


### OLS regression, bootstrap CI, depth vs mass

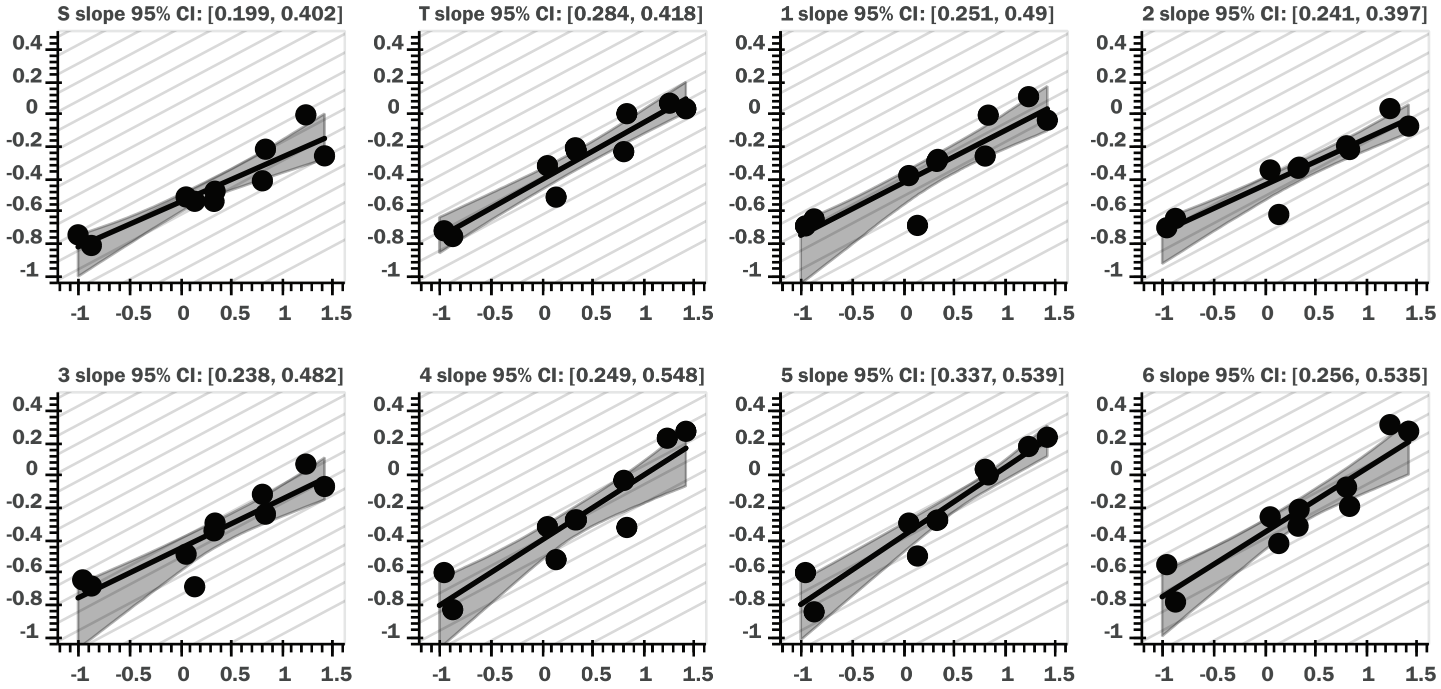

### Bayesian regression, area/depth vs mass

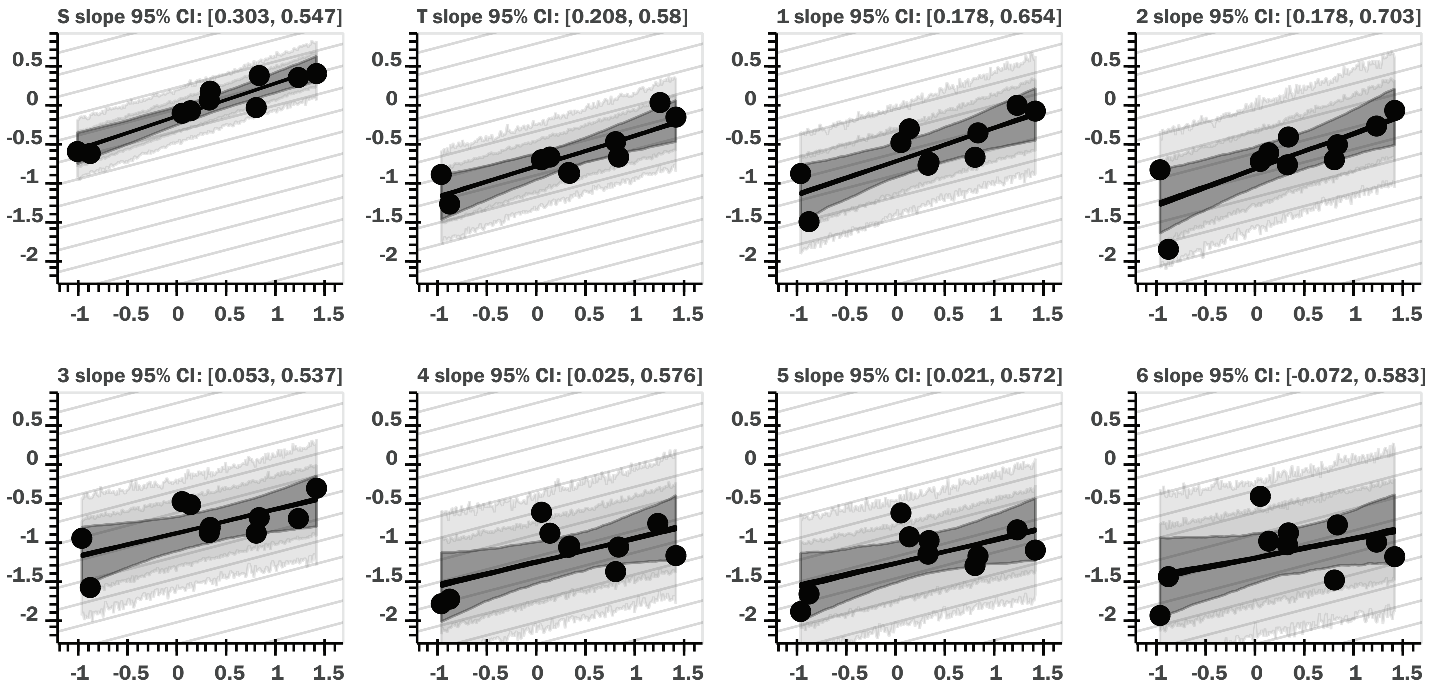


### OLS regression, bootstrap CI, area/depth vs mass

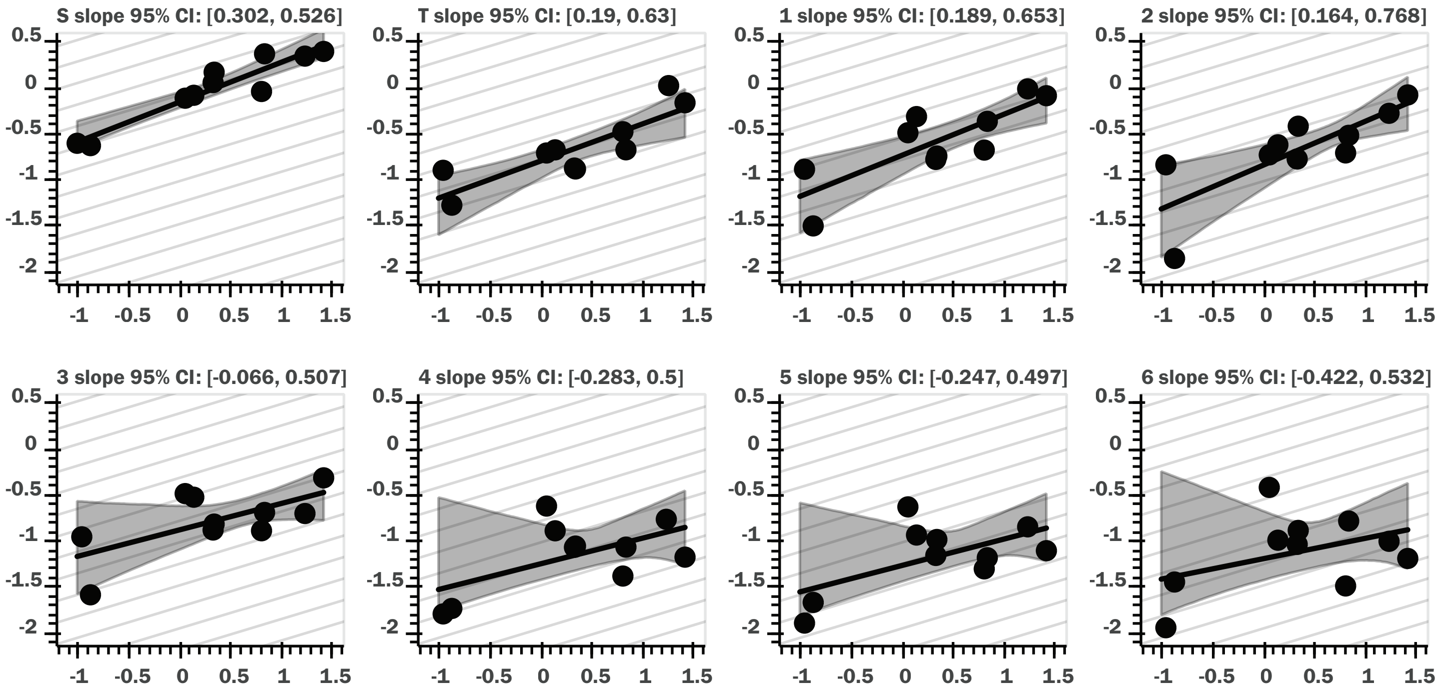

### Bayesian regression, area²/depth vs mass

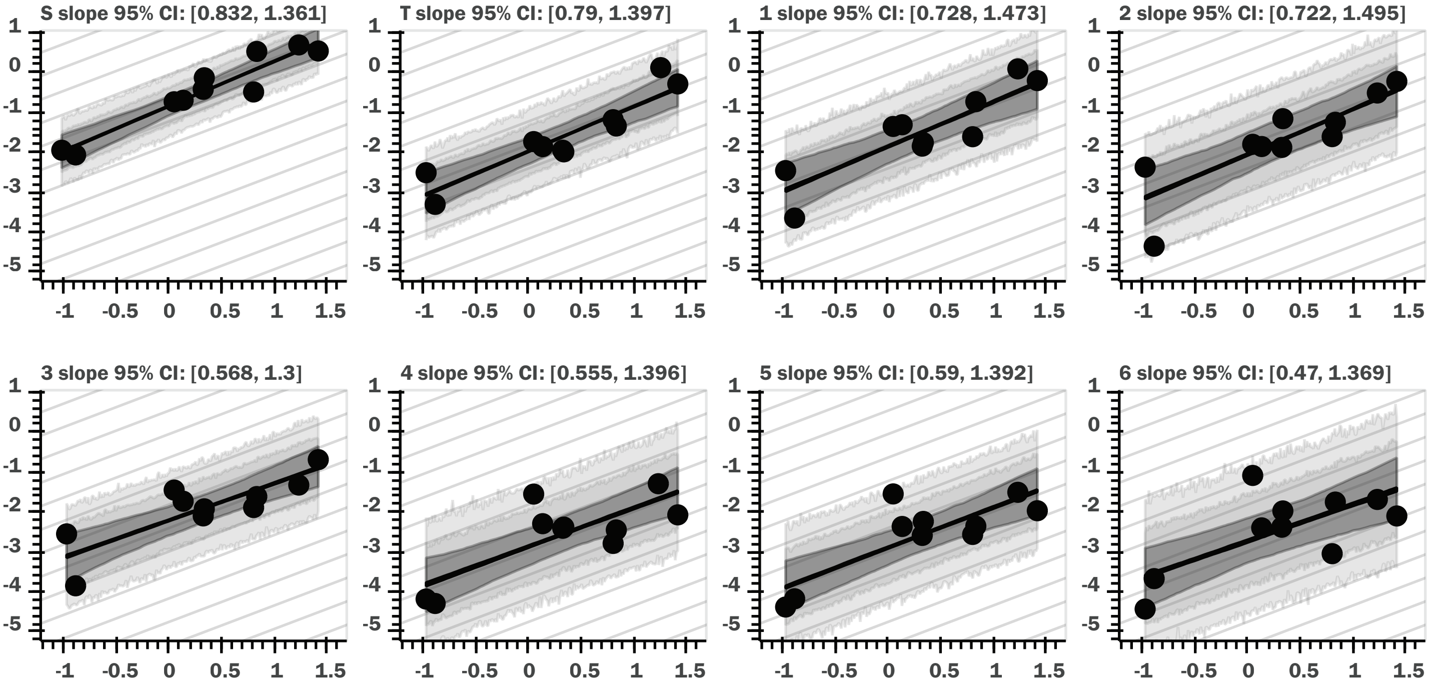


### OLS regression, bootstrap CI, area²/depth vs mass

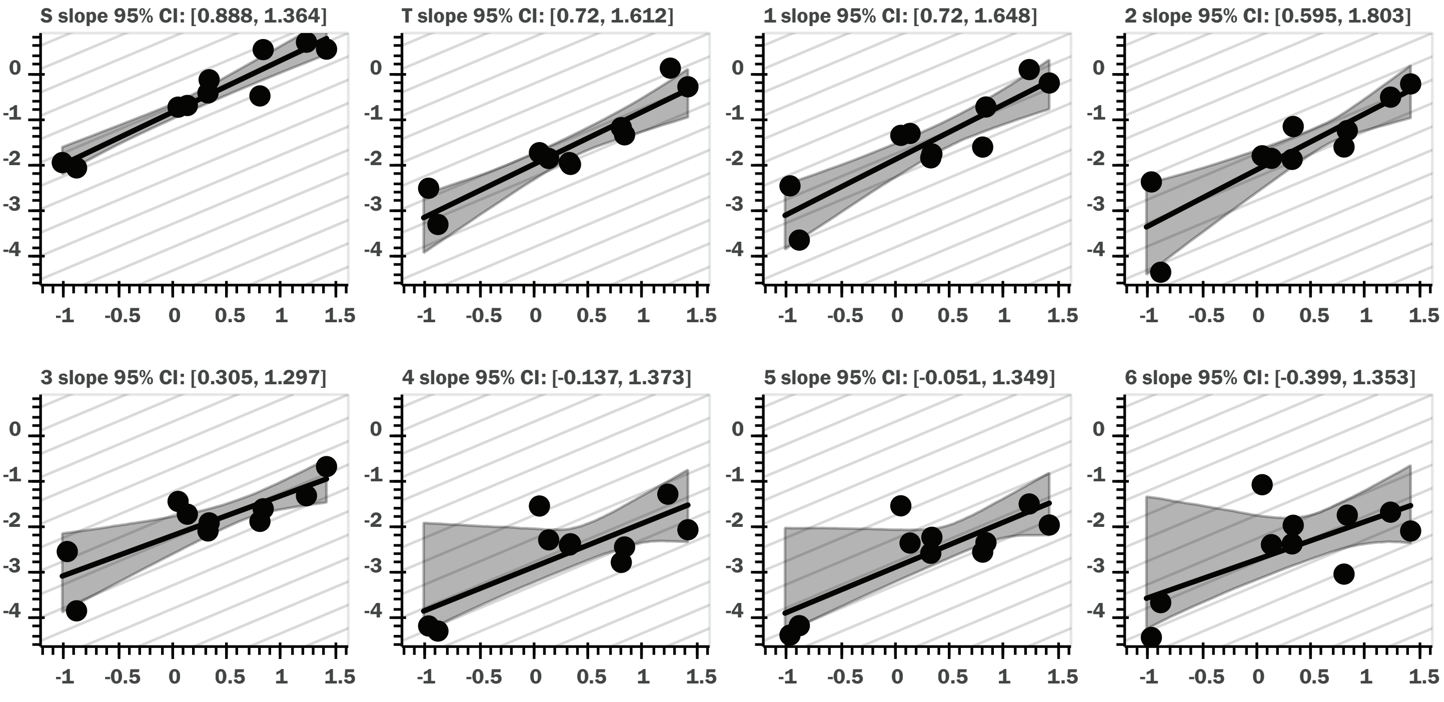
